# Vascular Basement Membrane Laminins Modulate Functional Zonation of Cerebral Microvessels

**DOI:** 10.1101/2025.11.17.688818

**Authors:** Tushar Deshpande, Kishan Kapupara, Melanie-Jane Hannocks, Jula Huppert, Sai-Kiran Samawar, Devika Rag Thulichery, Sigmund Budny, Sharang Ghavampour, Jian Song, Lingzhang Meng, Ralf H. Adams, Hyun-Woo Jeong, Lydia Wachsmuth, Cornelius Faber, Jens Soltwisch, Rupert Hallmann, Lydia Sorokin

**Author notes:** Max Planck Institute for Biological Intelligence, Martinsried, Germany (Jula Huppert). Department of Clinical Research and Epidemiology, German Centre for Cardiac Disease, University Clinic Wuerzburg, 97078 Wuerzburg, Germany (Sharang Ghavampour). Institute of Cardiovascular Sciences, Guangxi Academy of Medical Sciences, & The People’s Hospital of Guangxi Zhuang Autonomous Region, Nanning, Guangxi, China (Lingzhang Meng). These authors contributed equally to this work (Tushar Deshpande, Kishan Kapupara). Corresponding author: Lydia Sorokin, Institute of Physiological Chemistry and Pathobiochemistry, University of Muenster, Waldeyerstrasse 15, 48149 Muenster, Germany.

## Abstract

We investigated whether vascular basement membrane (BM) laminins influence vascular zonation by performing single-cell RNA sequencing on cerebral blood vessels from mice lacking the major vascular laminins in endothelial and smooth muscle BMs, laminin α4 (*Lama4^-/-^*) and laminin α5 (*Tek-cre:Lama5^-/-^*), and wild-type littermates. Our dataset expands existing cerebral vascular transcriptomic profiles and reveals that *Lama4^-/-^* endothelial cells exhibit increased arterial marker expression and reduced postcapillary venule identity. *In vitro* and *in vivo* studies indicate that compensatory upregulation of laminin α5 in *Lama4^-/-^*vessels enhances expression of junctional proteins (*Ocln*, *Cldn5*) and promotes vessel contractility via increased expression of contractile molecules in mural cells. Additionally, loss of *Lama4* upregulates expression of large artery markers (*Gja4*, *Dll4*, *Tgfb2*) and results in elevated autotaxin (*Enpp2*) levels, a key enzyme in lysophosphatidic acid production implicated in stroke. Accordingly, *Lama4^-/-^* mice exhibit worsened stroke outcomes, driven not by immune infiltration or junctional defects, but by increased vascular permeability likely mediated by autotaxin and/or activation of resident myeloid cells. Our data suggest that laminin α4/α5 ratios in vascular BMs regulate functional zonation between arterioles, capillaries and postcapillary venules by modulating metabolic pathways in endothelial and mural cells, and indirectly influencing resident myeloid cells.

## Introduction

Vascular zonation refers to spatial differences in structure, function, and molecular characteristics along the arteriovenous axis within an organ ^1^. In the brain, vascular zonation shapes blood-brain barrier (BBB) integrity, nutrient and metabolite exchange, and leukocyte entry points during neuroinflammation - factors that contribute to stroke severity and dementia risk.

Arterial identity is broadly established by flow and VEGF-induced Notch signalling, while venous identity is driven by COUP-TFII, which represses arterial gene expression (reviewed in ^2^). Although transitions between arteries, capillaries, and venules are associated with distinct morphological features and changes in blood flow and pressure, the molecular basis of these transitions remains incompletely understood. At the capillary level, vessel specialization (e.g., fenestrated vs. non-fenestrated, tight junction complexity) is shaped by organ-specific demands and the local cellular and extracellular environment. Advanced *in vivo* imaging has shown that mural cells - including smooth muscle cells and pericytes – also exhibit zone-specific morphologies and expression patterns of PDGFRβ and smooth muscle actin (SMA) ^3^, implicating them in the functional zonation of the vasculature ^4^.

In the brain, the neurovascular unit (NVU) - a tightly interconnected system of blood vessels and neural cells - is essential for maintaining central nervous system (CNS) homeostasis ^5^. The NVU includes endothelial and perivascular cells, which vary along the vascular tree, as well as astrocytes, neurons, and microglia, all engaged in continuous cross-talk. Recent studies have also identified long-term resident macrophages and fibroblasts located perivascularly, particularly around arteries, suggesting they may further contribute to vascular-neural communication and functional zonation of vessels ^6, 7^. All NVU components are embedded within specialized extracellular matrices (ECMs), with basement membranes (BMs) being the most abundant due to the brain’s dense vasculature ^8^. These BMs form structural and functional barriers between endothelial and mural cells, as well as between the vasculature and the CNS parenchyma, defining perivascular spaces that harbour long-lived myeloid and fibroblastic cells ^8–10^.

Like all BMs, those of cerebral blood vessels consist of a collagen type IV and laminin network, interconnected by proteoglycans, such as agrin and perlecan, and nidogens ^9^. Both collagen IV and laminin are multi-isoform families, whose specific combinations create biochemically and functionally distinct BMs across different organs and vascular segments ^9^. Laminins are heterotrimers composed of α, β and γ chains. In endothelial BMs, laminin-411 (α4β1γ1) and laminin-511 (α5β1γ1) are variably expressed along the vascular tree, forming a scaffold beneath endothelial cells and surrounding pericytes. Laminin α4 and α5 chains are also present in vascular smooth muscle cell BMs, typically in combination with laminin β2 and γ1 ^11^. Beyond the vessel wall, the endothelial tube with its associated pericytes and perivascular cells is further encased by an astroglial layer and a parenchymal BM, which demarcates the boundary with the brain parenchyma and contains laminins 211 and 111 ^8^.

The molecular heterogeneity of vascular and parenchymal BMs in both development and adulthood suggests roles beyond structural support or barrier formation. In particular, laminin α4, the first laminin expressed in the vascular system and broadly distributed, has been implicated in vascular tube formation and branching ^12–14^. By contrast, laminin α5 expression begins perinatally and predominates in arterial endothelial and smooth muscle BMs ^8^, with much lower levels in veins. Its expression decreases at postcapillary venules - the primary sites of leukocyte extravasation ^15–17^. Laminin α5 also stabilizes VE-cadherin at endothelial junctions, a process essential for resistance artery responses to shear stress ^18^. Together, these findings indicate that distinct laminin isoforms play specialized roles along the vascular tree, particularly within the microvasculature.

We here investigate whether vascular laminins contribute to the functional zonation of cerebral vessels by performing single-cell RNA sequencing (scRNA-seq) of cortical blood vessels from mice lacking laminin α4 (*Lama4^-/-^*) or laminin α5 (*Tek-cre:Lama5^-/-^*), major vascular BM components, and wild-type littermates. In *Lama4^-/-^*mice, compensatory upregulation of laminin α5 occurs ^13, 15^. *In vitro* and *in vivo* analyses showed that laminin α5 enhances the expression of junctional proteins (*Ocln*, *Cldn5*), suppresses markers of postcapillary venules and endothelial activation (*Vcam1*, *Icam1*), and promotes contractile gene expression in mural cells - altogether contributing to an artery-like phenotype. Conversely, laminin α4 suppresses large artery endothelial markers (*Gja4*, *Dll4*, *Edn1*, *Tgfb2*), supports the quiescent state of resident myeloid cells, and represses *Enpp2* (autotaxin), a key enzyme in lysophosphatidic acid (LPA) production implicated in stroke pathology ^19, 20^. Consistent with this, *Lama4^-/-^* mice exhibited more severe outcomes after transient middle cerebral artery occlusion (tMCAO) compared to wild-type and an endothelial-specific *Lama5* knockout (*Tek-cre:Lama5^-/-^*), a phenotype independent of immune cell infiltration and instead linked to elevated autotaxin and potentially also activated resident myeloid cells.

Together, our data indicate that the laminin α4/α5 balance in vascular BMs influences cerebral vessel zonation - particularly at arteriole-capillary and capillary-postcapillary venule transitions - by modulating endothelial and mural cell metabolism and indirectly affecting resident immune cell states.

## Material and Methods

### Mice

Age- (8-14 weeks) and sex-matched *Lama4^-/-^* ^13^ and *Tek-Cre:Lama5^-/-^*^21^ mice were used. As similar results were obtained for *Lama4^+/+^*and *Lama5^floxed/floxed^* mice, data were pooled and referred to as wild-type (WT) controls. Animal experiments were approved by the Landesamt für Verbraucherschutz und Ernährung, Nordrhein-Westfalen (LAVE; license numbers: 81-02.04.2023.A130 and 84-02.05.50.17.004), carried out according to German Animal Welfare Act guidelines and reported according to ARRIVE guidelines.

### Single cell RNA sequencing

Mice were transcardially perfused with ice-cold PBS, brains removed, minced in ice cold DMEM (Sigma-Aldrich) and digested for 60 min at 37°C in 30 U/ml papain (LK003153; Worthington Biochemical), 40 µg/ml DNase type IV (LS006331; Worthington Biochemical) and 0.125 mg/ml Liberase TM (5401119001; Sigma-Aldrich). Myelin was removed by 10 min 1000 x *g* centrifugation through 22% BSA. Brains from 3 WT and 3 *Lama4^-/-^* mice were pooled/sample.

Single cells were encapsulated in Gel Bead Emulsion using the 10X Chromium system (10X Genomics). Libraries were prepared according to manufacturer’s instructions using Chromium Single Cell 3’ Library & Gel Bead Kit v3 (10X Genomics) and sequenced in Illumina NextSeq 500 using High Output Kit v2.5 (150 cycles, Illumina).

The Cellranger (10x genomics) pipeline was followed to demultiplex the sequencing data, to align it to the mouse reference genome (mm10), to count the reads, and to aggregate the resulting cell-gene matrices according to sample replicates. Subsequent data analysis was performed with R package Seurat (version 3.1). Raw counts were normalized with SCTransform ^22^ and samples were integrated (reciprocal PCA) to correct for the batch effect ^23^. Dimensional reduction, unsupervised clustering, visualization and differential expression analysis was performed using Seurat. Gene set enrichment analysis (GSEA) was performed using GSEA and Molecular signature database (MSigDB) ^24^. RNA-seq data generated for this paper have been deposited in the NCBI GEO (GSE308748).

### Immunofluorescence

Brain preparations and immunofluorescence staining were as previously described ^8^. Primary and secondary antibodies employed are in Supplementary Tables 1 and 2. Sections were analysed using Zeiss confocal laser scanning (LSM 900 and LSM800) and Spinning-disk confocal microscopes. Images were analysed using Volocity 6.3 (Perkin-Elmer), Zen Blue (Zeiss) and Imaris 10 (Oxford Instruments). Mean fluorescence intensity was quantified in ZEN Blue from maximum intensity projections after applying a manually adjusted threshold to eliminate background.

Automated pericyte morphology and coverage quantification was performed in ZEN Blue. After Gaussian smoothing and automatic 3-sigma thresholding of raw images, pericyte and vessel areas were segmented to calculate pericyte coverage on the vascular surface. Outliers were manually curated post-batch analysis.

For 3D reconstruction of pericytes, vessel surfaces were rendered in Imaris 10 and pericytes were manually traced and color-coded based on morphology.

### Western blots

Brains were homogenized in RIPA buffer with protease inhibitors. Lysates (50 μg protein/lane) were separated under reducing conditions, except for PECAM-1 which was analysed under non-reducing conditions. After transfer to nitrocellulose, membranes were probed with antibodies listed in Supplementary Tables 1 and 2. Band intensities were quantified using ImageJ; due to lower vessel density in *Lama4^-/-^* brains, signal intensities of VCAM-1, claudin-5, prostacyclin synthetase and SMA were normalised to the PECAM-1 signal; autotaxin signal intensity was normalised to tubulin. GAPDH and tubulin served as loading controls.

### Flow cytometry

Mice were transcardially perfused with ice-cold PBS, brains removed, minced in ice cold DMEM and digested at 37°C for 60 min using papain (100 Units/brain) and DNase (0.1 mg/ml). Myelin was removed by 10 min 1000 x *g* centrifugation through isotonic Percoll (Amersham). Endothelial cells were stained with the antibodies listed in Supplementary Table 1 and analysed using a FACSCelesta (Becton Dickinson) and FlowJo.

### Transient middle cerebral artery occlusion (tMCAO)

Transient focal ischemia using the intraluminal filament model was performed on 12-week male mice ^25^. The left middle cerebral artery was occluded for 60 min and reperfused for 24h; ipsilateral (IL) brain hemispheres represent stroke samples, contralateral (CL) hemispheres represent non-stroke controls. Motoric scores were performed anonymously as described ^26^. Mice exhibiting no deficit (score 0) or with rolling to the paretic side (score 4) were excluded.

<u>For analysis of leukocytes after stroke,</u> IL and CL hemispheres were dissociated, subjected to a discontinuous density gradient centrifugation using Percoll and total cells were counted. Leukocytes were stained with the antibodies listed in Supplementary Table 1 and acquired using a FACSCalibur (Becton Dickinson) and analysed using FlowJo.

### Infarct volume measurement

Infarct volumes were measured using 5 x 1mm coronal sections incubated in 2% 2,3,5-Triphenyltetrazolium chloride (TTC) for 20 min at 37°C. Sections were fixed in 2% PFA and photographed. Infarct volume (*V*) was calculated as follows:

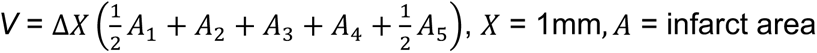

Swelling (S) of IL hemispheres was calculated and expressed as percent of the CL hemisphere:

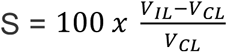, where *V_IL_* and *V_CL_* = total IL and CL volumes, respectively.

The edema-corrected infarct volume (Vi) was: 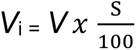

### Bulk RNA sequencing

Mice were perfused with PBS, and brains were isolated. After removing the olfactory bulb and cerebellum, the remaining IL and CL hemispheres were homogenized in TRIzol (Life Technologies). Brains were pooled (2/group). Differential centrifugation was performed to remove myelin and RNA was extracted via chloroform/phenol extraction and purified with DNase and RNeasy columns (Qiagen) for bulk RNA sequencing (Illumina NextSeq500).

FASTQ files were generated using bcl2fastq (version v2.17.1.14 Illumina). Subsequent analysis was performed using the Galaxy program. Quality control was performed using FASTQC (version 0.72+galaxy1). The last 10 low-quality bases were trimmed away. The trimmed reads (70 bp) were mapped with RNA STAR (version 2.6.0-b1) to the mouse reference genome (mm10) with gencode vM22 gene annotation. Reads were counted with featureCounts tool (version 1.6.4). The raw counts were normalized using DESeq2 package in R program. Differentially expressed genes were detected using unadjusted *p*-value (Wald’s test) and log2 fold change. Functional analysis was performed using GSEA. Normalized bulk RNA-seq data (GSE309181) was deconvoluted using SCADEN (Version 0.9.1) using existing single cell profiles ^27^.

### LPA measurements

MALDI-2-MS imaging was performed on a modified timsTOF fleX mass spectrometer (Bruker) ^28^. A sublimated 2,5 dihydroxyacetophenone matrix was used to acquire positive-ion mode images (200-2000 m/z, 25 µm pixel size). Data was processed in SCiLS Lab (Bruker), with lipid assignments based on accurate mass (5 ppm window).

### Statistics

Analyses used GraphPad Prism 9. Tests (specified in legends) were selected based on normality (assessed by Q-Q plots): parametric for normal data, non-parametric otherwise. For two groups, t-tests or Mann-Whitney tests were applied; multiple groups were compared by one-way ANOVA. Significance was defined as *p* < 0.05. Data are means ± SD.

## Results

### Single cell RNA sequencing of Lama4^-/-^ cerebral vessels

To comprehensively investigate differences between *Lama4^-/-^* and littermate WT cerebral blood vessels, single cell RNA sequencing (scRNA-seq) was performed on endothelial and associated mural and immune cell enriched samples isolated from the cerebral cortex of adult male animals; neurons were largely excluded by depleting myelin positive populations ^29^ (sFig. 1A). 18,828 single cell transcriptomes were clustered and annotated based on previous studies ^30, 31^ (sFig. 1B). UMAP plots show the distribution of cell types identified, confirming high proportions of endothelial (47%), perivascular (15%) and myeloid cells including microglia (29%) (sFig. 1C), which did not differ significantly between WT and *Lama4^-/-^*samples. Additional clustering characteristics are shown in sFig. 1D,E. This revealed low expression of most laminins in mature WT tissues and the absence of ECM expression in immune cells (sFig. 1F,G). Supplementary File 1 and 2 contain the complete gene lists (GSE308748); full names of genes occur in Supplementary Table 3.

### Endothelial laminins affect expression of arterial markers

Analysis of gene profiles of endothelial and mural cells showed a similar trend, with no changes from WT in absolute numbers of cells in *Lama4^-/-^* samples but with a shift towards higher expression of genes that typically associate with arterial endothelium (*Gja4*, *Edn1*, *DII4*, *Tgfb2*, *Gskn3* and *Klf2*) (Fig. 1A-C; sFig. 2A,B) and more contractile (*Acta2, Cnn1, Tagln, Fbln5*) or vasoactive mural cells (*Ptgis*) (Fig. 1D-G; sFig. 2C,D). Endothelial clustering revealed 6 groups (Fig. 1A; sFig. 2A), with a slightly higher proportion of cells expressing a large artery gene profile (*Gja4*, *Tgfb2, Edn1*, *DII4*) and capillaries with high junctional tightness (*Cldn5, Ocln*) in *Lama4^-/-^* compared to WT (Fig. 1B), and a reduction in the proportion of cells expressing genes associated with postcapillary venules/activated endothelial cells (*Icam1, Vcam1*) (Fig. 1C). In addition, high expression of *Enpp2* was observed in *Lama4^-/-^* endothelial and mural cells (Fig. 1B), which encodes autotaxin, required for lysophosphatidic acid (LPA) generation, and has been implicated in stroke ^19, 32^. Tissue-resident macrophages (Fig 1H-J, sFig. 2E,F) were also reduced in *Lama4^-/-^* mice, but had higher proportions of MHCII^+^ and phagocytotic populations (*Lyve1^low^, Mrc1, Wfdc17, Cd68*) (Fig. 1J), despite absence of ECM expression by these cells (sFig. 1F,G). Additionally, high *Ttr* (8.45 fold) expression in *Lama4^-/-^* endothelium and mural cells was observed (Fig. 1B,E), which encodes transthyretin that binds T4 and T3 in the circulation. While the higher expression of *Enpp2* was confirmed using various methods (see below), *Ttr* data was variable and could not be confirmed by qPCR of isolated cerebral vessels or endothelial cell lines (not shown).

**Figure 1.**
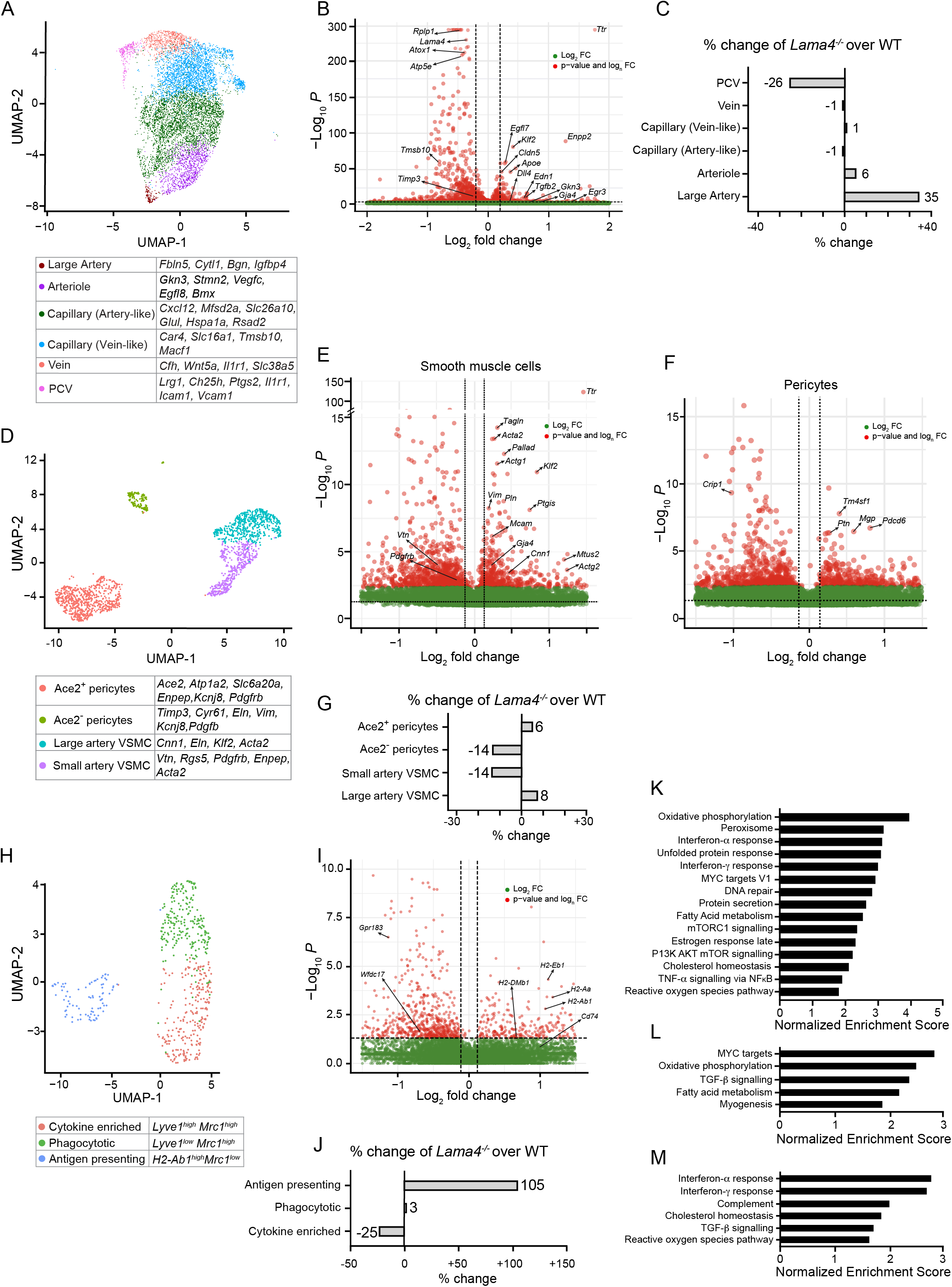
scRNA-seq data for endothelial cell populations (A-C), smooth muscle cells and pericytes (D-G) and resident immune cells (H-J) in enriched cerebral vessel samples from *Lama4^-/-^*and WT littermates. UMAPs identified **(A)** 6 endothelial **(D)** 2 smooth muscle and 2 pericyte, and **(H)** 3 myeloid cell populations. Different cells are colour coded based on their cluster affiliations shown in associated tables. Volcano plots of genes upregulated and downregulated in *Lama4^-/-^* samples compared to WT littermates in **(B)** endothelial cells, **(E)** smooth muscle cells **(F)** pericytes and **(I)** myeloid cells; **(C,G,J)** corresponding graphs of percentage changes in *Lama4^-/-^* over WT littermates based on transcriptomic profiles. Unbiased GSEA analyses showing enrichment of pathways in *Lama4^-/-^*samples compared to WT littermates in **(K)** endothelial cells, **(L)** all mural cells and **(M)** resident myeloid cells.

Unbiased gene set enrichment analysis (GSEA) revealed enrichment of metabolic gene sets, including oxidative phosphorylation and fatty acid metabolism, in *Lama4^-/-^*endothelial and mural cells (Fig. 1K,L), as well as enhanced proteolysis in *Lama4^-/-^*endothelial cells (Fig. 1K) and, in mural cells, genes associated with myogenesis and TGFβ signalling (Fig. 1L). Similar analyses for tissue-resident macrophages revealed enrichment of IFNα and IFNγ responses, cholesterol homeostasis and TGFβ signalling (Fig. 1M).

### Compensatory Lama5 upregulation enhances junctional tightness and mesh-like pericyte phenotype

As *Lama4^-/-^* mice show compensatory enhanced laminin α5 deposition in endothelial BMs ^15, 33^, scRNA-seq was performed on isolated cerebral vessels from mice lacking endothelial laminin α5 expression (*Tek-cre:Lama5^-/-^*) (sFig. 3). 63,050 cells were analysed, which showed the same clusters as for *Lama4^-/-^* analyses. This revealed the opposite pattern to that observed in *Lama4^-/-^* endothelium with regards to *Vcam1*, *Ocln and Klf2* (sFig. 3A-C), i.e. upregulation of the leukocyte adhesion molecule, *Vcam1* (and *Lama4),* and downregulation of the tight junction molecule *Ocln,* and the shear response gene *Klf2*, suggesting that some results observed in *Lama4^-/-^*endothelium are due to the elevated expression of laminin α5. Immunofluorescence staining confirmed more VCAM-1 positive vessels, in particular postcapillary venules, in *Tek-cre:Lama5^-/-^* and fewer in *Lama4^-/-^* brains (Fig. 2A). A similar pattern was found for ICAM-1, another leukocyte adhesion molecule and marker of postcapillary activation^34^ (sFig. 4A,B). Immunofluorescence staining for occludin revealed less intense staining at endothelial cell-cell junctions in *Tek-cre:Lama5^-/-^* brains and more intense staining in *Lama4^-/-^*brains compared to WTs (Fig. 2B). Staining for another tight junction molecule, claudin 5, revealed more microvessels showing high intensity staining in *Lama4^-/-^* brains and slightly fewer in *Tek-cre:Lama5^-/-^*brains compared to WTs (Fig. 2C). Western blots of brain extracts, normalized to PECAM-1^+^ endothelium, confirmed higher VCAM-1 on *Tek-cre:Lama5^-/-^* and slightly lower levels on *Lama4^-/-^* endothelium (Fig. 2D,E), and higher claudin 5 on *Lama4^-/-^* endothelium (Fig. 2D,F). In addition, qPCR showed higher expression of both *Ocln* and *Cldn5* in endothelial cells sorted from *Lama4^-/-^* brains and lower expression in those from *Tek-cre:Lama5^-/-^* brains compared to controls (sFig. 4C-E). Plating of a brain endothelial cell line (bEND.5) on laminin 411 versus 511 *in vitro* showed higher *Cldn5* and *Ocln* expression on laminin 511 (sFig. 4F).

**Figure 2.**
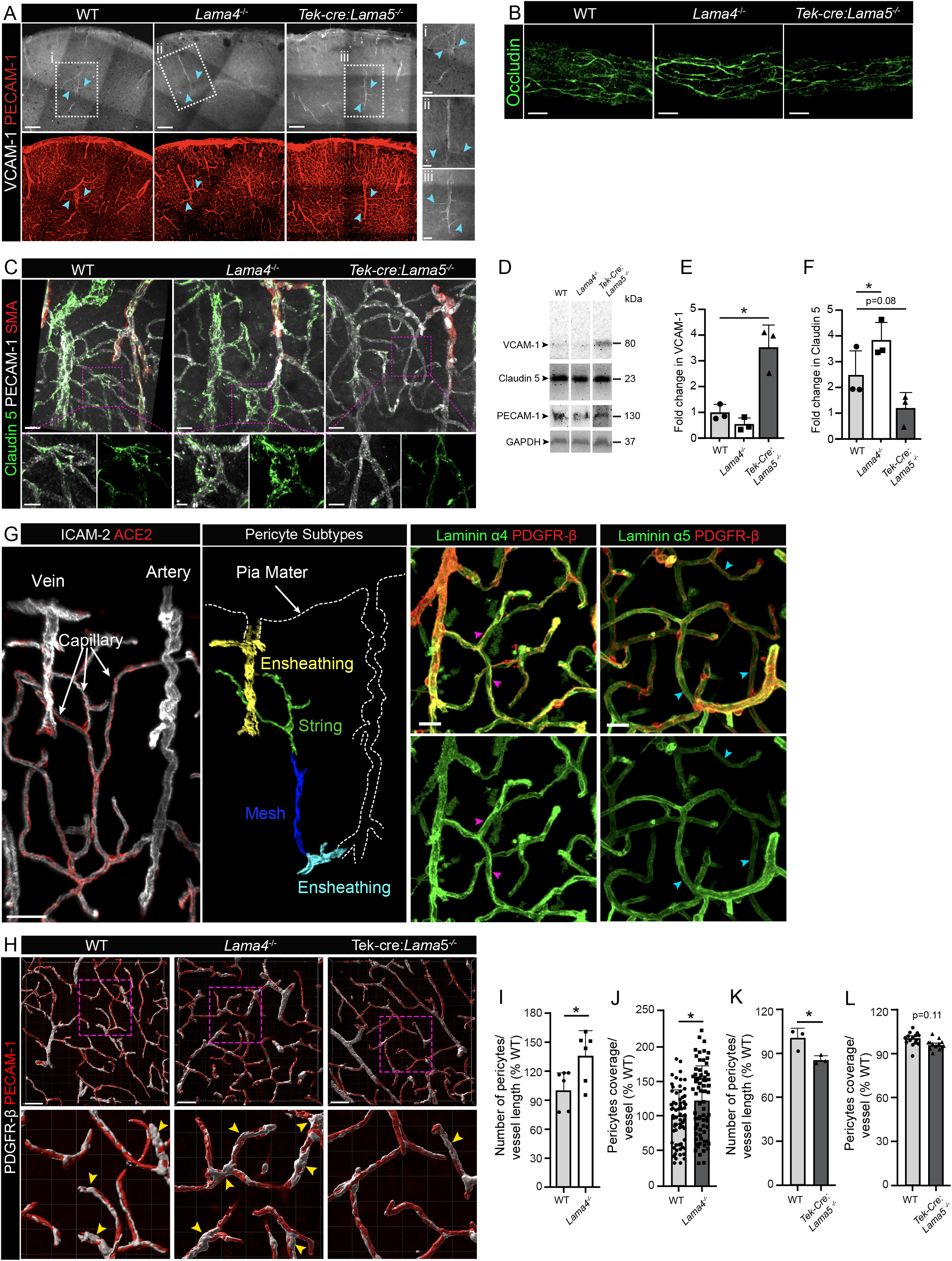
Enhanced endothelial junctional molecule expression and more mesh-like pericytes in *Lama4*^-/-^ vessels due to compensatory upregulation of *Lama5*. **(A)** Immunofluorescence staining of *Lama4^-/-^*, *Tek-Cre:Lama5*^-/-^ and representative WT control brain sections for VCAM-1. Higher magnifications are shown to the right. Immunofluorescence staining of *Lama4^-/-^*, *Tek-Cre:Lama5*^-/-^ and representative WT control brain sections for (**B**) occludin and **(C**) claudin 5. Higher magnifications show claudin 5 expression on capillary endothelial cells. **(D)** Representative Western blots for VCAM-1, Claudin 5 and PECAM-1 in *Lama4^-/-^*, *Tek-Cre:Lama5*^-/-^ and WT brain extracts and (**E, F**) corresponding quantification of signal intensities normalized to PECAM-1 (one-way ANOVA, 3-5 mice per genotype in at least 2 separate experiments). **(G)** Immunofluorescence staining for ACE2^+^ pericytes and ICAM-2^+^ endothelial cells identified ensheathing, string and mesh pericytes along the vascular tree. Immunofluorescence staining for PDGFRβ^+^ pericytes together with laminin α4 or laminin α5 in WT brain sections shows the presence of laminin α4 (magenta arrowheads) and lack of laminin α5 (cyan arrowheads) in string and mesh pericytes in representative WT control brain sections. (**H**) PDGFRβ^+^ string and mesh pericytes on PECAM-1^+^ vessels in WT, *Lama4^-/-^*and *Tek-Cre:Lama5*^-/-^ mice (yellow arrowheads indicate mesh pericytes in the higher magnifications). Quantification of PDGFRβ^+^ pericyte number and coverage per vessel length in *Lama4^-/-^* **(I, J)** and *Tek-Cre:Lama5*^-/-^ **(K, L)** brains expressed as a percentage of the appropriate WT control; (Unpaired t-Test, 3-5 mice per genotype in at least 2 separate experiments, in **J** and **L** each data point indicates an image analysed); data are means ± SD. \**p*<0.05, \*\**p*<0.01, \*\*\**p*<0.001. Scale bars: (**A**) 200 µm, (**B**) 10 µm, (**C**) 20 µm, (**C** higher magnification) 10 µm, (**G**) 30 µm and (**H**) 40 µm.

Analysis of lung endothelial cells isolated from transgenic mice showed a similar pattern of results, indicating that laminin α5 induced enhanced expression of junctional molecules is not specific for the brain (sFig. 4C-E). As VCAM-1 and ICAM-1 are endothelial activation markers, expressed by postcapillary venules ^34^, these data suggest that laminin α5 in the endothelial BM suppresses activation and that *Lama4^-/-^*mice have a reduced proportion of vessels that function as postcapillary venules (Fig. 1C), while *Tek-cre:Lama5^-/-^* mice have more (sFig. 3C). This is consistent with the reduced extravasation observed in *Lama4^-/-^* mice and enhanced extravasation in *Tek-cre:Lama5^-/-^* mice in different inflammatory models ^15, 33, 35^.

Interestingly, *Tek-cre:Lama5^-/-^*vessels also showed the opposite gene expression profile to that observed in *Lama4^-/-^* pericytes (sFig. 3D-F), which are embedded within the endothelial BM, suggesting that some of the effects observed in *Lama4^-/-^* mural cells, in particular pericytes, result from compensatory enhanced expression of laminin α5, rather than loss of laminin α4. Staining for pericytes using ACE2 allowed identification of the 3 types of pericytes along the vascular tree: ensheathing, string and mesh (Fig. 2G) ^36^. Staining for PDGFRβ together with laminins α4 or α5, revealed intense laminin α4 staining along pericyte processes of string and mesh pericytes, which was not the case for laminin α5 (Fig. 2G), consistent with the higher *Lama4* expression in pericytes relative to endothelium in scRNA-seq analyses of WT mice (sFig. 1F). PDGFRβ staining and quantification of pericyte cell bodies per blood vessel length revealed increased pericyte numbers and coverage in *Lama4^-/-^* brains (Fig. 2H-J), with high magnification revealing a more mesh-like phenotype ^4^ (Fig. 2H). The opposite results were found in *Tek-cre:Lama5^-/-^* vessels (Fig. 2H, K, L), which showed a 13% reduction in pericyte number (Fig. 2K) and a 5% reduction in pericyte coverage (Fig. 2L), reflecting a decrease in mesh-like phenotype and an increase in string pericytes. Mesh pericytes are smooth muscle actin negative; their branching processes allow them to interact with endothelial cells and other pericytes, potentially facilitating communication and coordination of blood flow regulation ^4^. This is consistent with the elevated *Klf2* expression in endothelium and mural cells and the exacerbated shear response in *Lama4^-/-^* mice ^18^, and the reduced *Klf2* expression pattern (sFig. 3E) and almost complete absence of shear response in *Tek-cre:Lama5^-/-^* mice ^18^.

No difference was noted in *Enpp2 (*autotaxin), *Gja4, Edn1 or Tgfb2* expression in either endothelium or mural cells of *Tek-cre:Lama5^-/-^* mice (sFig. 3B), nor were there major differences in resident myeloid populations (sFig. 3G,H), indicating that these changes in *Lama4^-/-^* where due to the loss of laminin α4.

### Laminin α4 suppresses some arterial characteristics

*In vivo* immunofluorescence staining for smooth muscle actin (SMA) and PECAM-1 was used to distinguish arteries from veins, revealing fewer and slightly shorter arteries in *Lama4^-/-^* brains (Fig. 3A-C) but with a 15% larger diameter compared to WT controls (Fig. 3D); no significant differences were observed for *Tek-cre:Lama5^-/-^*arteries (not shown). This is in accordance with previous data showing outward remodelling of *Lama4^-/-^* resistance mesenteric arteries, probably due to chronic elevated shear stress ^18^. Although the SMA^+^ vessels were shorter, staining intensities per vessel were higher in *Lama4^-/-^* compared to WT brains (Fig. 3E). Quantification of the number of SMC cells per vessel length revealed similar results for *Lama4^-/-^* and WT vessels, confirming higher expression of SMA in *Lama4^-/-^* VSMCs (Fig. 3F,G) as suggested by the scRNA-seq data. In addition, *Ptgis* was one of the most highly upregulated genes in *Lama4^-/-^*smooth muscle cells (Fig. 1E), which encodes prostacyclin synthetase required for prostacyclin synthesis, a potent vasoactive lipid mediator that leads to vasodilation. Western blots of *Lama4^-/-^* brain extracts confirmed higher amounts of SMA and prostacyclin synthetase (Fig. 3H-J). The increased prostacyclin synthetase expression probably contributes to the exacerbated shear-induced vasodilation in *Lama4^-/-^* mice _18._

**Figure 3.**
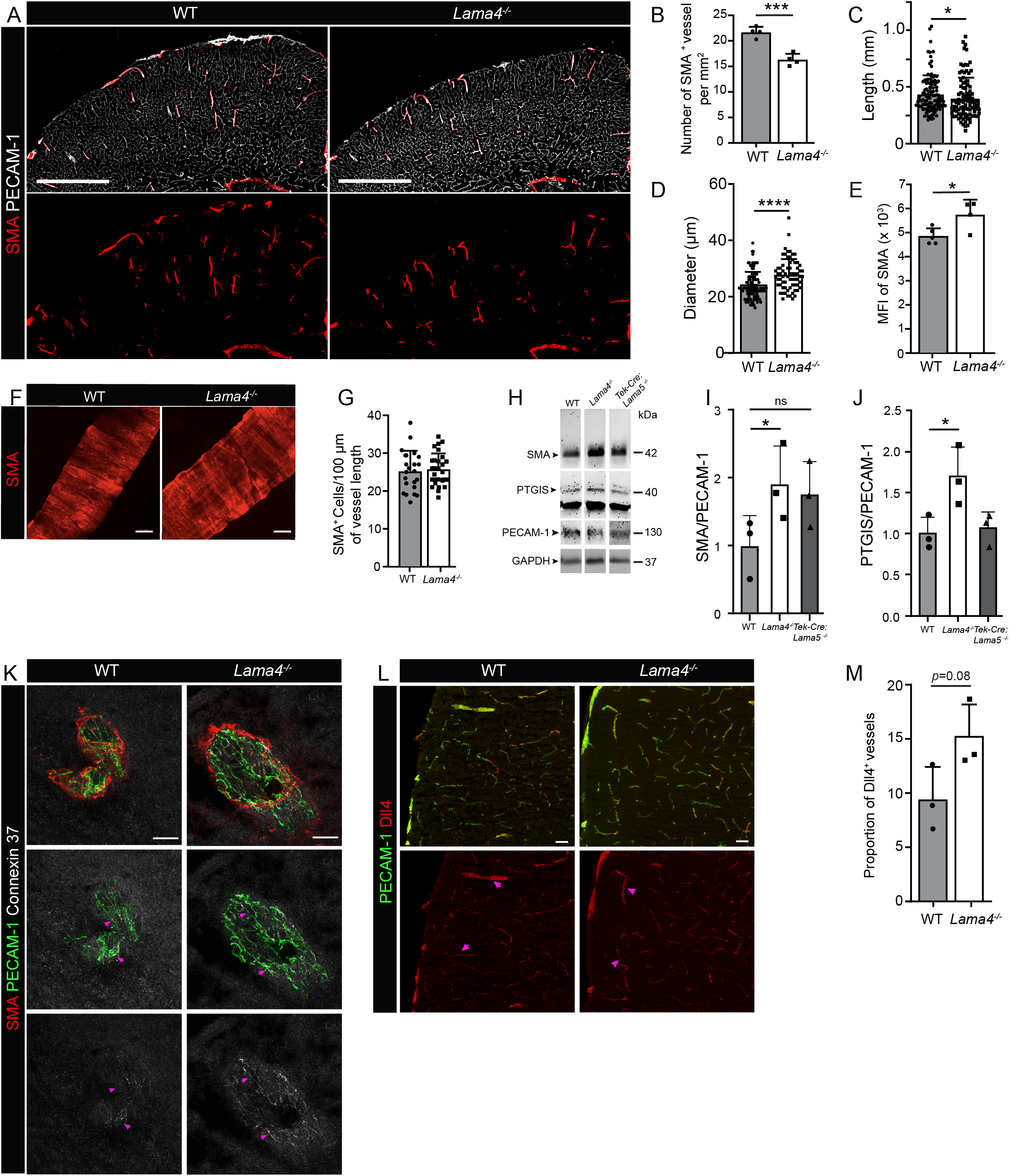
Laminin α4 suppresses some arterial endothelial markers. **(A)** Immunofluorescence staining of PECAM-1 and smooth muscle actin (SMA) to mark arteries in *Lama4^-/-^* brains and corresponding quantification of **(B)** number, (**C**) length (**D**) diameter and (**E**) mean fluorescence intensity (MFI) of SMA^+^ penetrating arteries (unpaired t-Test for **B**, **D**, **E** and Mann-Whitney for **C**; at least two 200 µm sections from 3-5 brains per genotype were analysed in at least 2 experiments; in **C** and **D**, each data point represents an artery). **(F)** High resolution image of SMA^+^ arteries in *Lama4^-/-^* and WT littermate brain sections and (**G**) corresponding quantification of SMA^+^ cells per vessel length (unpaired t-Test, at least two 200 µm sections from 3 brains per genotype were analysed in at least 2 experiments, each data point represents an artery). **(H)** Representative Western blots for smooth muscle actin (SMA) and prostacyclin synthetase (PTGIS) in *Lama4^-/-^*, *Tek-Cre:Lama5*^-/-^ and WT brain extracts with (**I, J**) corresponding quantification of signal intensities normalized to PECAM-1 (one-way ANOVA, 3-5 mice per genotype in at least 2 separate experiments). (**K**) Immunofluorescence staining for connexin 37 at arterial endothelial junctions (magenta arrowheads) and (**L**) of Dll4 (magenta arrowheads) in *Lama4^-/-^* and WT littermate brain sections and (**M**) corresponding quantification of Dll4^+^ vessels expressed as a percent of total vessels counted (unpaired t-Test, at least 3 sections from 3 brains per genotype were analysed in at least 2 experiments). Data are means ± SD. \**p*<0.05, \*\**p*<0.01, \*\*\**p*<0.001. Scale bars: (**A**) 500 µm, (**F**) 10 µm (**K**) 25 µm and (**L**) 50 µm.

Stainings for connexin 37 (encoded by *Gja4*) in arteries showed enhanced expression levels at endothelial junctions (Fig. 3K), while staining for delta-like 4 (*Dll4*), a marker of arterial endothelium, showed slightly higher intensity and broader staining (10%) in *Lama4^-/-^* arteries (Fig. 3L, M).

To investigate potential differences in tissue perfusion resulting from differences in vessel size or density, magnetic resonance imaging (MRI) was performed. The basal cerebral blood flow (CBF), as measured by arterial spin labelling, was not different between WT and *Lama4^-/-^* mice at the time points checked (6, 8 and 14 weeks) (sFig. 5A). The microvessel density index, however, was lower in the *Lama4^-/-^*mice (sFig. 5B), suggesting a slight reduction in vessel density as observed in stainings for ICAM-2 (sFig. 5C,D) and by flow cytometry (sFig. 5E). *Tek-cre:Lama5^-/-^*mice showed no reduction in vessel density as shown by flow cytometry (sFig. 5F).

Hence, there is a slightly lower vessel density in *Lama4^-/-^* brains but more vessels exhibit arterial characteristics and perfusion of the organ is normal.

### Indirect effects of loss of vascular Lama4 expression

Despite the absence of expression of laminins by immune cells (sFig. 1F,G), resident myeloid populations associated with the cerebral arteries and the pia ^37^ showed clear differences between WT and *Lama4^-/-^* brains (Fig. 4A). Macrophages defined by *Mrc1* (which encodes CD206)*, Ms4a7, Pf4* expression separated into 3 clusters (Fig. 1H; sFig. 2E,F), as previously reported ^7^; however, with some differences. We identified macrophages enriched in phagocytosis markers (*Mrc1, Wfdc17, Cd68, Ctsd*) with low *Lyve1* expression, a cytokine enriched population (*Ccl7, Ccl3 Atf3, Ednrb*) exhibiting high *Lyve1* expression, and a Lyve1 negative population characterised by high expression of markers of antigen presentation (*H2-Ab1* which encodes MHCII) and leukocyte infiltration (*Ccr2*) (Fig. 1I,J). We refer to these macrophage populations as phagocytotic, cytokine-enriched and MHCII^high^ antigen presenting populations, respectively. *Lama4^-/-^*mice had higher proportions of the MHCII^high^ (*H2-Ab1*) population, which were not restricted to the choroid plexus as reported by others ^7^ but occurred both perivascularly and in the meninges as shown by immunofluorescence staining (Fig. 4A). Immunofluorescence staining for CD206, MHCII and pan-laminin revealed an overall reduction in perivascular macrophages and confirmed that a higher proportion of CD206^+^ macrophages were MCHII^high^ in *Lama4^-/-^* brains (Fig. 4A); this was confirmed by flow cytometry of brains (without choroid plexus) (Fig. 4B). Hence, although there were fewer resident macrophages, they had an activated phenotype.

**Figure 4.**
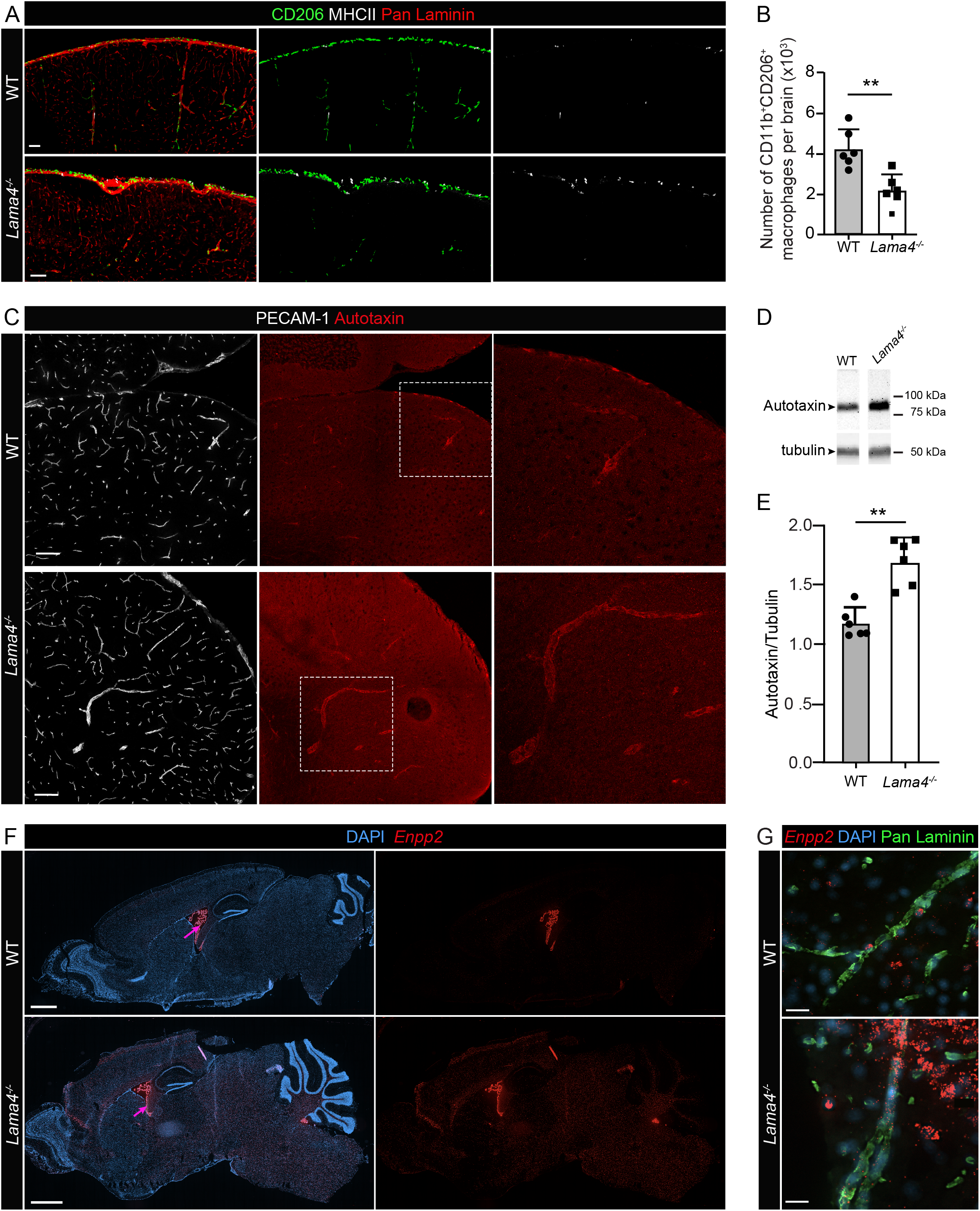
Indirect effects of loss of *Lama4* expression on resident myeloid populations and autotaxin expression. **(A)** Triple immunofluorescence staining for CD206^+^ resident macrophages, MHCII and pan laminin to mark vascular and pial BMs in *Lama4^-/-^*and WT littermates brain sections. **(B)** Flow cytometry quantification of CD11b^+^CD206^+^ macrophages in *Lama4^-/-^*or WT littermate brains (unpaired t-Test, 6 brains per genotype were analysed in at least 2 experiments). **(C)** Double immunofluorescence staining for PECAM-1 to mark blood vessels and autotaxin in *Lama4^-/-^* and WT littermates brain sections; right panels show boxed areas at higher magnification. **(D)** Representative Western blot for autotaxin in brain extracts from *Lama4^-/-^* and WT mice and **(E)** corresponding quantification of signal intensity normalized to the tubulin control (Unpaired t-Test, 6 mice per genotype in at least 2 separate experiments). **(F,G**) RNAscope together with immunofluorescence staining for DAPI as a nuclear marker and pan laminin to mark vascular BMs. Data are means ± SD \**p*<0.05, \*\**p*<0.01, \*\*\**p*<0.001. Scale bars: in (**A**) 100 µm, (**C**) 100 µm, (**F**) 1 mm and (**G**) 10 µm.

Although *Ttr* was highly expressed in all *Lama4^-/-^*vascular cells analysed, *in vitro* confirmation of this result using qPCR of isolated cerebral vascular cells, staining of tissue sections or Western blots for transthyretin in brain sections were inconclusive due to high variability. By contrast, elevated expression of autotaxin encoded by *Enpp2* in *Lama4^-/-^* brains was confirmed by immunofluorescence staining and Western blots of brain extracts (Fig. 4C-E). RNAscope revealed *Enpp2* expression mainly by perivascular cells, probably pericytes and fibroblasts, and by choroid plexus epithelium in WT brains, as previously reported ^38^, and enhanced expression in vascular and brain parenchymal cells in *Lama4^-/-^*brains (Fig. 4F, G). Furthermore, immunofluorescence staining and RNAscope both suggested enhanced expression of autotaxin protein (Fig. 4G) by brain parenchymal cells (sFig. 1F). This suggests that the elevated autotaxin expression in *Lama4^-/-^*brains may in part be due to indirect effects, such as the activated status of resident myeloid cells or prolonged enhanced shear stress, both of which promote autotaxin expression ^39, 40^.

### tMCAO as a test of arterial function

To further test arterial functions in *Lama4^-/-^* mice and to assess the functional consequence of elevated autotaxin, which is known to worsen stroke outcome ^20, 32^, a transient middle cerebral artery occlusion (tMCAO) model was employed. 60 min tMCAO and 24h reperfusion was performed in *Lama4^-/-^* and age- and sex-matched WT controls and in *Tek-Cre:Lama5^-/-^*mice as a control for effects specifically due to loss of laminin α4. *Lama4*^-/-^ mice exhibited more severe stroke symptoms as assessed by motoric activity, swelling and lesion volume compared to WT littermates and *Tek-Cre:Lama5*^-/-^ mice (Fig. 5A-C). However, flow cytometry revealed similar numbers of CD45^+^ leukocytes in the ipsilateral (IL) brain hemispheres in *Lama4*^-/-^, *Tek-Cre:Lama5*^-/-^ and WT mice (Fig. 5D), and similar ratios of total CD45^+^, neutrophils, macrophages, monocytes or microglia in IL compared to contralateral (CL) hemispheres (sFig. 6A), confirming our earlier observations that stroke severity at early stages after ischemia does not correlate with leukocyte extravasation into the parenchymal tissue ^25^. Immunofluorescence staining of brain sections for pan-laminin and CD45 revealed most leukocytes within or closely associated with arterioles located close to the surface of the brain in all mice (Fig. 5E, sFig. 6B), as previously shown for WT mice ^25^. The enhanced stroke severity observed in *Lama4*^-/-^ mice, therefore, was not determined by the extent of immune cell infiltration, nor can it be explained by junctional differences given the lack of a stroke phenotype in *Tek-cre:Lama5^-/-^* mice but is consistent with the higher autotaxin levels in these mice (Fig. 4C-E).

**Figure 5.**
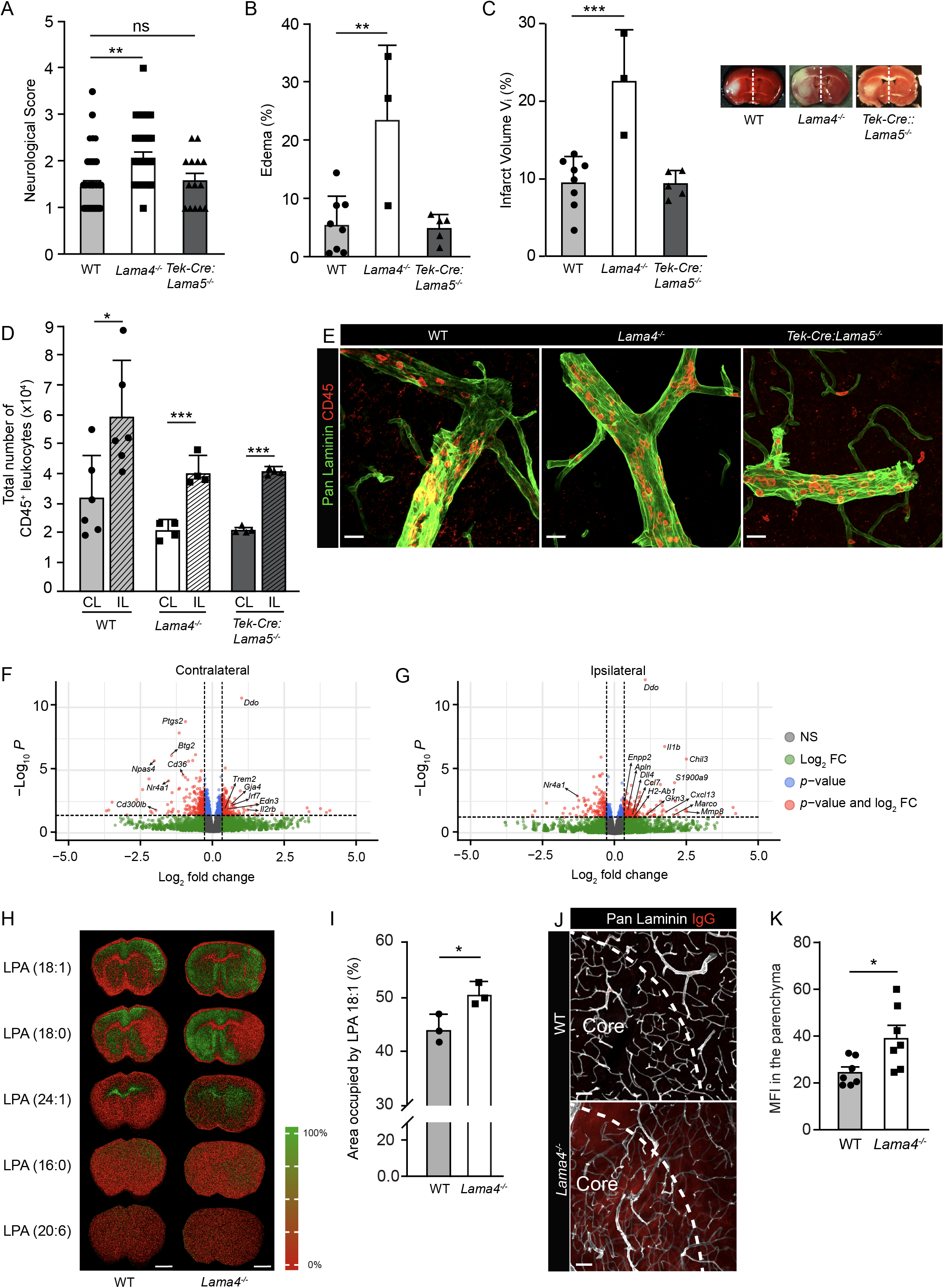
More severe tMCAO (60min/24h reperfusion) in *Lama4*^-/-^ correlates with higher LPA levels. (**A**) Neurological scores, (**B**) swelling in ipsilateral (IL) hemispheres and (**C**) infarct volumes in *Lama4^-/-^*, *Tek-Cre:Lama5*^-/-^ and WT controls (ANOVA followed by post hoc Tukey; for **A** at least 14 mice per genotype and, for **B** and **C**, at least 3 brains per genotype in at least 3 experiments)). (**D**) Flow cytometry quantification of total CD45^+^ leukocytes in IL and contralateral (CL) hemispheres of *Lama4^-/-^*, *Tek-Cre:Lama5*^-/-^ and WT controls (ANOVA followed by post hoc Tukey, 3-6 brains per genotype from at least 3 experiments). (**E**) Representative immunofluorescent staining for pan-laminin to mark the border to the CNS parenchyma and CD45^+^ leukocytes in brain sections from *Lama4^-/-^*, *Tek-cre:Lama5^-/-^* and WT controls. (**F**) Volcano plots showing relative differences in expression levels of all differentially expressed genes (DEGs) between WT and *Lama4^-/-^* mice for CL (F) and IL (**G**) hemispheres (total number of genes analysed = 20346); *p* < 0.05 and –0.7 ≥ Log_2_ fold change ≤ 1.3 are shown in red (DEGs = 204 (contralateral) and 213 (ipsilateral)). (**H**) Representative mass spectrometry imaging visualizing the signal intensity distribution of m/z: 437.27 tentatively assigned to LPA in the post stroke brain. Oedema enlarged hemisphere is the ipsilateral side. LPA (18:1) is increased on the ipsilateral side relative to the contralateral in both *Lama4^-/-^* and WT mice. (**I**) % Area occupied by LPA (18:1) signal (unpaired t-Test, 3 mice per genotype in at least 2 experiments). **(J)** Representative immunofluorescent staining for pan-laminin and IgG in *Lama4^-/-^* and WT brain sections and **(K)** corresponding quantification of IgG immunofluorescent signal in the CNS parenchyma (unpaired t-Test, 7 brains per genotype were analysed in 3 experiments). Data are means ± SD \**p*<0.05, \*\**p*<0.01, \*\*\**p*<0.001. Scale bars: (**E**) 25 µm, (**H**) 2 mm and (**J**) 60 µm.

### Bulk RNA Sequencing

Bulk RNA sequencing performed on the cerebral cortex from IL and CL brain hemispheres from *Lama4*^-/-^ and WT controls at 24h after ischemic stroke, confirmed the higher degree of neuron damage in *Lama4*^-/-^ mice and demonstrated that stroke-induced transcriptomic differences dominated over differences due to genotype (sFig. 7A, B). Volcano plots show fold changes in gene expression and *p* values in CL and IL (Fig. 5F) brain hemispheres from *Lama4*^-/-^ compared to WT mice. Supplementary Files 3 and 4 contain the complete gene lists (GSE309181).

Despite the prominent stroke effects, comparison of IL hemispheres from WT and *Lama4*^-/-^ mice also revealed upregulation of several arterial markers in *Lama4*^-/-^ including *Gja4*, *Dll4*, *Gkn3*, *Edn1,* as well as inflammatory parameters such as *Il1b* (interleukin-1b), *Ccl7*, *Mmp8*, *Chil3*, *Cxcl13* and *Marco,* and *Enpp2* (autotaxin) (Fig. 5F,G), substantiating the scRNA-seq data. The heatmap in sFig. 7C shows the top genes using a cut-off of *p* ≤ 0.05, listed according to fold-change. GSEA revealed enrichment of metabolic gene sets (oxidative phosphorylation), ribosomal genes, and gene sets involved in muscle in CL *Lama4*^-/-^ brain hemispheres (sFig. 7D), consistent with the scRNA-seq data. In IL *Lama4*^-/-^ brain hemispheres, GSEA showed enrichment for metabolic processes, cytokine signalling, Toll-like receptor and NOD-like receptor signalling (sFig. 7E), suggesting a proinflammatory status. Given the absence of enhanced infiltration of immune cells in the IL hemisphere of *Lama4^-/-^* mice compared to WT, changes in cytokine signalling and Toll-like receptor genes are likely to represent differences in resident myeloid populations.

*Enpp2* encodes autotaxin, a phosphodiesterase that converts lysophosphatidylcholine (LPC) to lysophosphatidic acid (LPA), a lipid mediator that acts through several G-coupled protein receptors (LPAR1-6) with diverse effects including ones on vascular integrity ^41, 42^. As LPA levels exacerbate stroke severity ^20^, we employed *in situ* mass spectrometry to analyse the LPA species in IL and CL hemispheres of *Lama4^-/-^* and WT brains after tMCAO, revealing elevated levels of several LPA species (18:1, 18:0, 24:1) in *Lama4^-/-^* brains compared to WT controls, with enrichment in the IL hemispheres (Fig. 5H,I) ^32^.

As autotaxin affects vessel permeability ^43^, the presence of serum proteins, including IgG, in the CNS parenchyma of *Lama4^-/-^* mice was assessed using immunofluorescence staining, revealing higher intensity and broader staining in the lesion core and penumbra (Fig. 5J, K). To determine whether blood vessels of *Lama4*^-/-^ mice are generally more permeable than those of WT or *Tek-Cre:Lama5^-/-^* mice, a modified Miles permeability assay was performed ^44^. Extravasation of intravenously injected Evan’s blue dye in response to intradermal injection of PBS into the back skin of age- and sex-matched mice revealed no differences between *Lama4*^-/-^, *Tek-Cre:Lama5*^-/-^ or WT mice (sFig. 8), indicating similar basal permeabilities. However, an approximately two-fold higher extravasation of Evans Blue into the dermis of *Lama4*^-/-^ mice was measured following intradermal injection of the vascular endothelial cell permeability factor (VEGF) compared to *Tek-Cre:Lama5*^-/-^ or WT mice (sFig. 8), indicating enhanced permeability to soluble molecules but only under stress conditions, which is consistent with the generally higher levels of autotaxin in these mice.

The more severe stroke phenotype in the *Lama4^-/-^* mice compared to *Tek-cre:Lama5^-/-^* mice is consistent with the elevated levels of autotaxin (and consequently LPA), given that this was the major difference between these two phenotypes. However, the fact that autotaxin is upregulated in several cell types in the brain parenchyma of *Lama4^-/-^*mice, which do not express laminin α4, suggests that the proinflammatory phenotype, resulting from changes in resident myeloid populations or exacerbated shear response probably contribute to elevated autotaxin and higher stroke severity in these mice.

## Discussion

Our data demonstrate that BM laminins exert segment-specific effects along the vascular tree. Notably, *Lama4^-/-^* mice exhibited reduced vascular density and a shift toward an arterial phenotype, primarily at the expense of postcapillary venule identity. This was accompanied by increased expression of the shear-responsive gene *Klf2*, elevated levels of contractile and vasoactive molecules in smooth muscle cells, and a higher number of mesh-like pericytes per vessel ^4^. These observations suggest that the balance between laminin α4 and α5 in the microvascular BM plays a role in establishing the functional zonation of cerebral vessels. High levels of laminin α5 in endothelial BMs appear to promote an arterial phenotype, not only in endothelial cells but also in pericytes enclosed within the same BM. Conversely, laminin α4 suppresses the expression of key arterial markers, such as connexin 37, Delta-like 4 and TGF-β2, and is essential for maintaining the quiescent state of perivascular myeloid cells, which preferentially associate with arteries.

Elevated expression of autotaxin in *Lama4^-/-^* brains likely results from chronic shear stress - evidenced by outward arterial remodelling ^18^ - and/or activation of resident myeloid cells. This is consistent with worsened outcomes following transient middle cerebral artery occlusion (tMCAO) in *Lama4^-/-^* mice.

Gene expression profiling and immunofluorescence staining of *Lama4^-/-^* and *Tek-Cre:Lama5^-/-^* tissues, along with qPCR of sorted cells and *in vitro* studies using bEND.5 cells, support the conclusion that compensatory upregulation of laminin α5 in *Lama4^-/-^* vessels underlies the enhanced expression of junctional proteins (claudin-5 and occludin), consistent with our previous studies ^18, 45^, and suppresses the expression of the endothelial activation markers, ICAM-1 and VCAM-1. In *Lama4^-/-^*vessels, the higher number of pericytes and altered pericyte morphology are also likely due to this compensatory laminin α5 upregulation, given that *Tek-Cre:Lama5^-/-^* vessels exhibit the opposite phenotype. Specifically, the rise in mesh pericytes, which are α-smooth muscle actin-negative and associated with the arteriole-to-capillary transition ^4^, may enhance vascular permeability under VEGF stimulation and contribute to the exacerbated shear response in the *Lama4^-/-^* mice ^18^. These mesh pericytes have been implicated in blood flow regulation and blood-brain barrier maintenance ^3, 46^, and possess branching processes that facilitate interactions with endothelial cells and neighbouring pericytes, supporting coordinated vascular function. This is consistent with elevated *Klf2* expression in both endothelial and mural cells in *Lama4^-/-^* mice and aligns with this heightened shear sensitivity. Interestingly, increased pericyte coverage correlated with reduced VCAM-1 and ICAM-1 expression in *Lama4^-/-^* mice, as previously reported in other mouse strains ^47^, while *Tek-Cre:Lama5^-/-^* mice showed the opposite trend. This suggests that pericytes embedded in laminin α5-rich endothelial BMs may contribute to maintaining endothelial quiescence.

As expected, *Tek-Cre:Lama5^-/-^*vessels did not show significant changes in SMC gene expression or Western blots; however, a potential role of laminin α5 in enhancing contractile gene expression in SMCs in *Lama4^-/-^* mice cannot be excluded, and requires analysis of a vascular smooth muscle specific *Lama5* knockout mouse.

Importantly, elevated arterial marker expression (e.g., *Gja4, Dll4,* and *Tgfb2)* was observed exclusively in endothelial cells lacking laminin α4, supporting the notion that laminin α4 negatively affects arterial differentiation. Interestingly, the arterial expansion in *Lama4^-/-^* mice was at the expense of postcapillary venules, not venular endothelium *per se*. To our knowledge, this is the first report of a factor defining postcapillary venule identity that is not simply related to vessel diameter.

We have previously shown that laminin α5 expression begins in arterial vessels during embryonic development and increases postnatally ^48^. Laminins 411/511 modulate Notch1 signalling during retinal angiogenesis ^14^, and laminin α5 has been reported to inhibit proliferation of various cell types, including epithelial and hematopoietic cells ^49–51^. Our pathway analyses reveal a shift toward enhanced oxidative phosphorylation and lipid metabolism in *Lama4^-/-^* mice, consistent with an arterial phenotype and a quiescent state ^2^. Our data therefore suggest that vascular laminins do not determine arterial fate during development but rather maintain arterial identity - laminin α5 promotes, while laminin α4 inhibits this phenotype. This model is supported by the diminished shear responses in resistance arteries in *Tek-Cre:Lama5^-/-^* mice ^18^ - opposite to the *Lama4^-/-^* phenotype ^15, 18^.

Some of the changes observed on *Lama4^-/-^*vessels are likely to be indirect, e.g., the elevated endothelial expression of TGF-β2, which is known to upregulate expression of smooth muscle actin and other contractile proteins ^52–54^, may at least partially contribute to the enhanced contractile phenotype of SMCs in *Lama4^-/-^*vessels. Along the same lines, both bulk and scRNA-seq analyses revealed an inflammatory phenotype in *Lama4^-/-^*mice with clear increases in cytokine expression, and an associated reduction in resident macrophages localised along arteries, despite the fact that immune cells do not express ECM molecules, as confirmed here. These results suggest a role for vascular BMs in either the seeding and/or maintenance of perivascular myeloid populations, but potentially also their activation status, topics that are currently under investigation. It cannot be determined here whether changes in these resident perivascular macrophages also contributed to the more severe stroke in *Lama4^-/-^* mice.

Interestingly, elevated autotaxin expression in *Lama4^-/-^* brains by cells not normally expressing laminin α4 may result from chronic shear stress and/or inflammation as suggested by others ^55, 56^. Autotaxin has been implicated in various inflammatory conditions and modulates purinergic signaling ^57^. Elevated autotaxin levels are associated with worse stroke outcomes ^20, 32^, and endothelial-specific deletion of *Enpp2* (autotaxin) reduces stroke severity ^32^. The worsened stroke phenotype in *Lama4^-/-^*mice is consistent with these findings, and supports the hypothesis that laminin α4 loss indirectly drives autotaxin upregulation via inflammatory and potentially also biomechanical cues.

Finally, our scRNA-seq analysis extends existing profiles of cerebral vascular cells. We identified a likely postcapillary venule endothelial population (marked by *Icam1, Vcam1, Il1r1*) that was diminished in *Lama4^-/-^* mice. This may represent the activated endothelial subset previously described in WT EAE ^29^, potentially contributing to the reduced EAE severity observed in *Lama4^-/-^* mice ^15^. Furthermore, we distinguish large arteries and arterioles by smooth muscle gene expression, with high *Klf2* and *Acta2* (αSMA) levels in arteries versus *Vtn* and *Pdgfrb* in arterioles.

Together, these data indicate that the ratio of laminin α4 to α5 in vascular BMs regulates the functional zonation of cerebral vessels by influencing metabolic pathways in endothelial and mural cells and by modulating the activation and maintenance of resident myeloid populations. This finely tuned balance underlies the specialization of arterioles, capillaries, and postcapillary venules in the cerebral vasculature.

## Acknowledgements

We thank Dirk Reinhardt (EIMI) for stroke surgeries; Laura Schnell and Melissa Chmara for animal breeding, genotyping and tissue collection.

## Author contribution statement

TD performed the scRNA-seq and bulk RNA-seq analyses and verified data with immunofluorescence. KK verified a large part of the scRNA-seq and bulk RNA-seq data using flow cytometry, immunofluorescence and Western blots. SB contributed to Western blots. JH performed stroke experiments and bulk RNA seq. SG and LM contributed to the analyses of junctional proteins. S-KS and DRT analysed resident macrophage populations. M-JH contributed to verification of the scRNA-seq data using immunofluorescence and contributed to writing of the manuscript. JSong, RA and H-WJ contributed to planning and execution of the scRNA-seq experiments. LW and CF performed the MRI. JS performed the *in situ* mass spectrometry for LPA species. RH contributed to the project design and LS conceived the project, acquired funding and wrote the manuscript.

## Statements and Declarations

### Ethical considerations

Experiments on mice were approved by the Landesamt für Verbraucherschutz und Ernährung, Nordrhein-Westfalen (LAVE; animal permit license numbers: 2018.A068 and Histo4), carried out in accordance with the German Animal Welfare Act guidelines and reported according to ARRIVE (Animals in Research: Reporting *In Vivo* Experiments) guidelines.

### Consent to participate

Not applicable

### Consent for publication

Not applicable

## Supplementary Material

Supplementary material for this paper can be found at the journal website: (*link will be given by the journal)*

