## Supplementary data for "Vascular Basement Membrane Laminins Modulate Functional Zonation of Cerebral Microvessels"

[1]Institute of Physiological Chemistry and Pathobiochemistry and Cells-in-Motion Interfaculty Centre (CiMIC), University of Muenster, 48149 Muenster, Germany; [2]Department of Tissue Morphogenesis, Max-Planck-Institute for Molecular Biomedicine, 48149 Münster, Germany. [3]Clinic of Radiology, University of Muenster, 48149 Muenster, Germany, [5]Institute of Hygiene, University of Muenster, 48149 Muenster, Germany.

Current addresses: [†]Max Planck Institute for Biological Intelligence, Martinsried, Germany; [§]Department of Clinical Research and Epidemiology, German Centre for Cardiac Disease, University Clinic Wuerzburg, 97078 Wuerzburg, Germany; [#] Institute of Cardiovascular Sciences, Guangxi Academy of Medical Sciences, & The People's Hospital of Guangxi Zhuang Autonomous Region, Nanning, Guangxi, China.

*These authors contributed equally to this work.

Corresponding author:

Lydia Sorokin
Institute of Physiological Chemistry and Pathobiochemistry
University of Muenster, Waldeyerstrasse 15, 48149 Muenster, Germany

**Supplementary methods**

FACS sorting

Brains and lungs from mice were processed for sorting after transcardial perfusion with ice-cold PBS. Tissues were minced in ice-cold DMEM (Sigma-Aldrich) and digested at 37ºC for 1.5 hours using collagenase II (7.5 mg/ml; CLS2, Worthington Biochemical). Myelin was removed by centrifugation for 10 min at 1000 x *g* through 25% BSA (brain) or 0.5% BSA (lung). Endothelial cells were sorted using a FACSAria™ Fusion Cell Sorter (Becton Dickinson) using antibodies listed in Supplementary Table 1 and the strategy shown in Supplementary Figure 4. Data analysis was performed using FlowJo.

qPCR

Brains from 8-10 week-old male WT and *Lama4*$^{-/-}$ mice were prepared as above. Total RNA was extracted using a RNeasy Mini Kit (Qiagen). RNA was reverse transcribed to cDNA using the Omniscript RT Kit (Qiagen). Quantitative PCR was performed on a Rotor-Gene Q (Qiagen) using SYBR Green QPCR MasterMix (Agilent). Primers used are listed in Supplementary Table 4 (https://pga.mgh.harvard.edu/primerbank/). Gene expression analyses were performed using the ΔΔCT method [1] and normalised to the reference gene (*Gapdh*) values from the same sample. Results are expressed as percent increase/decrease in *Lama4*$^{-/-}$ compared to WT samples. At least 3 WT and 3 *Lama4*$^{-/-}$ mice in 4 separate experiments, with technical duplicates for each gene were evaluated.

Permeability experiments

The hair from the backs of mice was removed and 100 µl 1% Evans Blue dye was injected in the lateral tail vein; after 15 min either 50 µl of $VEGF_{165}$ (100 ng) or PBS was injected intradermally into two separate spots on the back skin (modified Miles assay). After 30 min, the mice were killed and a 0.5 $cm^2$ piece of skin was dissected from each area. The Evans Blue dye was eluted from the dissected skin by incubation with formamide at RT for 5 days. The amount of eluted dye was quantitated by spectrophotometry at 620 nm and the results expressed as $OD_{620}$/0.5 $cm^2$/5 days [2].

Magnetic resonance imaging (MRI)

To determine the microvessel density index (MDI) and cerebral blood flow (CBF), MRI was performed on 14 week-old naïve WT and *Lama4*$^{-/-}$ mice. Mice were anesthetized with isoflurane (5% induction, 1-1.5% maintenance) and imaged in a 9.4-T MRI (Bio-Spec 94/20; Bruker BioSpin MRI GmbH, Germany) equipped with a brain surface coil. Axial T2-weighted anatomical images were obtained: Repetition Time (TR) = 2000 ms, Echo Time (TE) = 50 ms, Rapid Acquisition with Relaxation Enhancement (RARE) factor = 8, 4 averages, field of view = 1.6 cm, matrix = 128 x 128, slice thickness = 1.5 mm, in plane resolution = 125 x 125 $\mu m^2$. Perfusion was assessed with a 1 slice arterial spin labelling (ASL) experiment with a Flow-sensitive Alternating Inversion Recovery-RARE protocol; field of view (FOV) $(16\ mm)^2$, slice thickness 1.5 mm, 1 slice, TR/Effective TE = 18000/44.8 ms, 22 inversion recovery times (TIR) 26.8-5000 ms, inversion slab thickness = 4.5 mm. Vessel parameters (blood volume, microvessel density) were determined by acquiring apparent diffusion coefficient (ADC) maps and ΔT2 and ΔT2* maps before and after i.v. administration of superparamagnetic 20 nm iron oxide particles (Ferucarbotran) at a dose of 30 mg Fe/kg [3]. Multi Slice Multi Echo (MSME) images for T2 mapping and multiple gradient echo (MGE) MR images for T2* mapping were obtained with the same geometry (FOV $(16\ mm)^2$, matrix $64^2$, slice thickness 0.3 mm). MSME was acquired with TR=5000 ms and 10 echos, TE = 10.9, 21.8, 32.7, 43.6, 54.5, 65.4, 76.3, 87.2, 98.1,109 ms. MGE was acquired with TR=1400 ms and 10 echos, TE = 4, 8,12,16,20,24,28, 32 ms with a 60° hermite pulse. The post contrast image acquisition was delayed by 3 min. Data for an ADC map with the same geometry were additionally acquired before contrast agent application, with a diffusion-weighted Echo Planar Imaging protocol (TR/TE 7500/18.4 ms) with b = 0, 300, 800 $s/mm^2$. Total scan time was approximately 45 min per animal.

For MRI data analysis, a Matlab (R2010b) routine was used for defining the region of interest (ROI) and calculation of ΔR2, ΔR2* and ADC maps. The cortical ROI in each data set was manually segmented across consecutive slices in the pre-contrast MSME image at TE=10.9 ms and was used to manually define the ROI across each of the 10 images of the time series. Microvessel density index (MDI) was determined as described in [3]. Fiji (Image J) and used to calculate cerebral blood flow from arterial spin labelling images.

### Supplementary Figures

Suppl. Figure 1

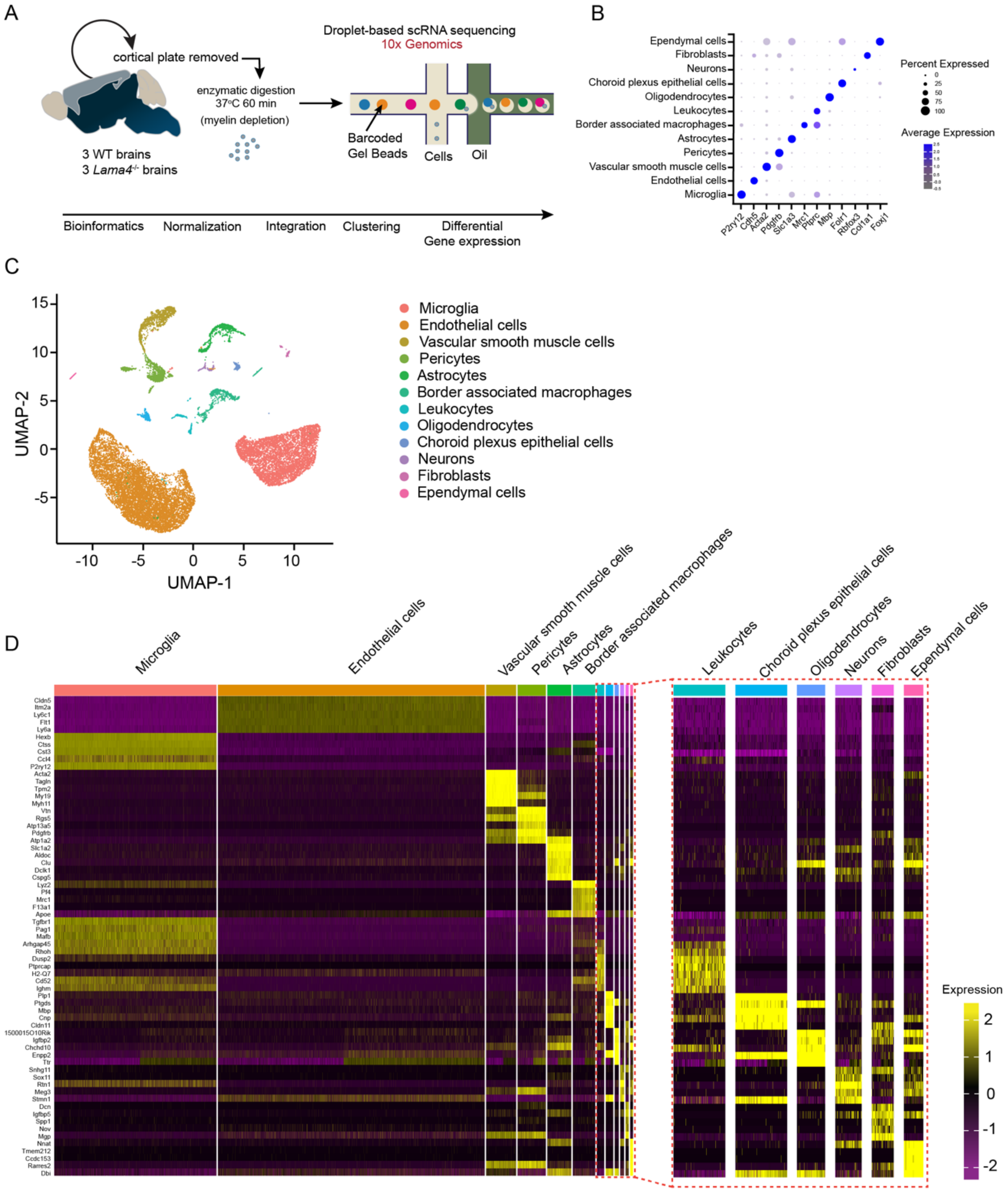

Suppl. Figure 1 continued

E

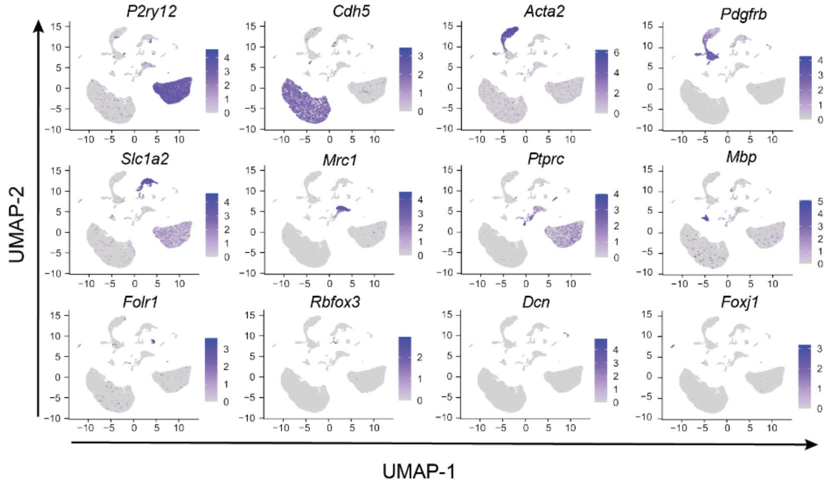


F

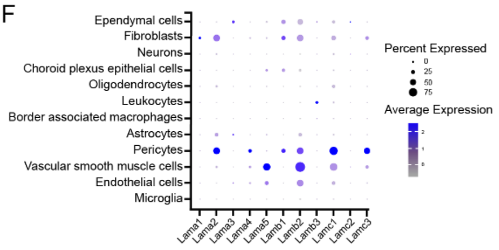


G

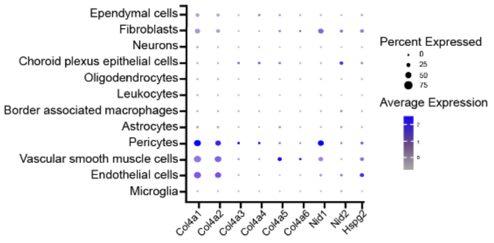


**sFigure 1 Single cell RNA sequencing data from neuron depleted samples from naïve *Lama4*$^{-/-}$ and WT control brains.**

(**A**) Schematic representation of the 10X chromium scRNAseq procedure used to deplete neurons and enrich blood vessel associated cells. (**B**) Dot plot showing major cell type markers. (**C**) Uniform Manifold Approximation and Projection (UMAP) of 18 828 cells from 6 samples (3 WT, 3 *Lama4*$^{-/-}$) depicting the different cell populations in the brain; each dot represents a cell; cells are colour coded based on their cluster affiliations. (**D**) Heatmaps representing the top 5 marker genes for each cluster. (**E**) UMAP feature plots showing the expression of key marker genes expressed by different cell populations. (**F**) Dot plot showing relative expression of different laminin isoforms for the different cell populations. (**G**) Dot plot showing relative expression of other basement membrane molecules for the different cell populations.

Suppl. Figure 2

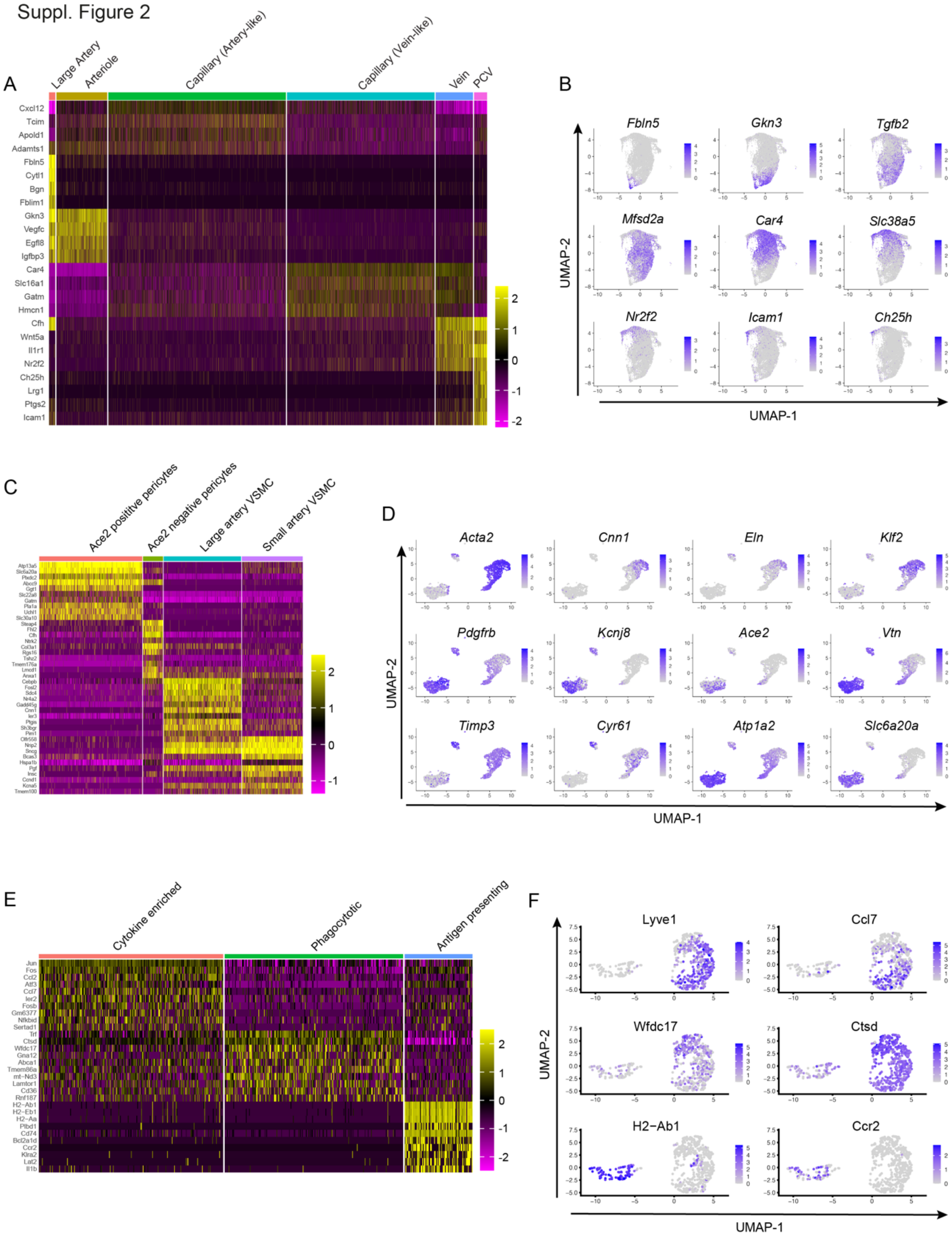


**sFigure 2 Single cell RNA sequencing data from neuron depleted samples from naïve *Lama4*$^{-/-}$ and WT control brains.** Heatmaps representing the top 5 marker genes for the different clusters in (**A**) endothelial cells, (**C**) mural cells and (**E**) myeloid cells. Corresponding UMAP feature plots showing the expression of key marker genes expressed by different cell populations in (**B**) endothelial cells, (**D**) mural cells and (**F**) myeloid cells.

Suppl. Figure 3

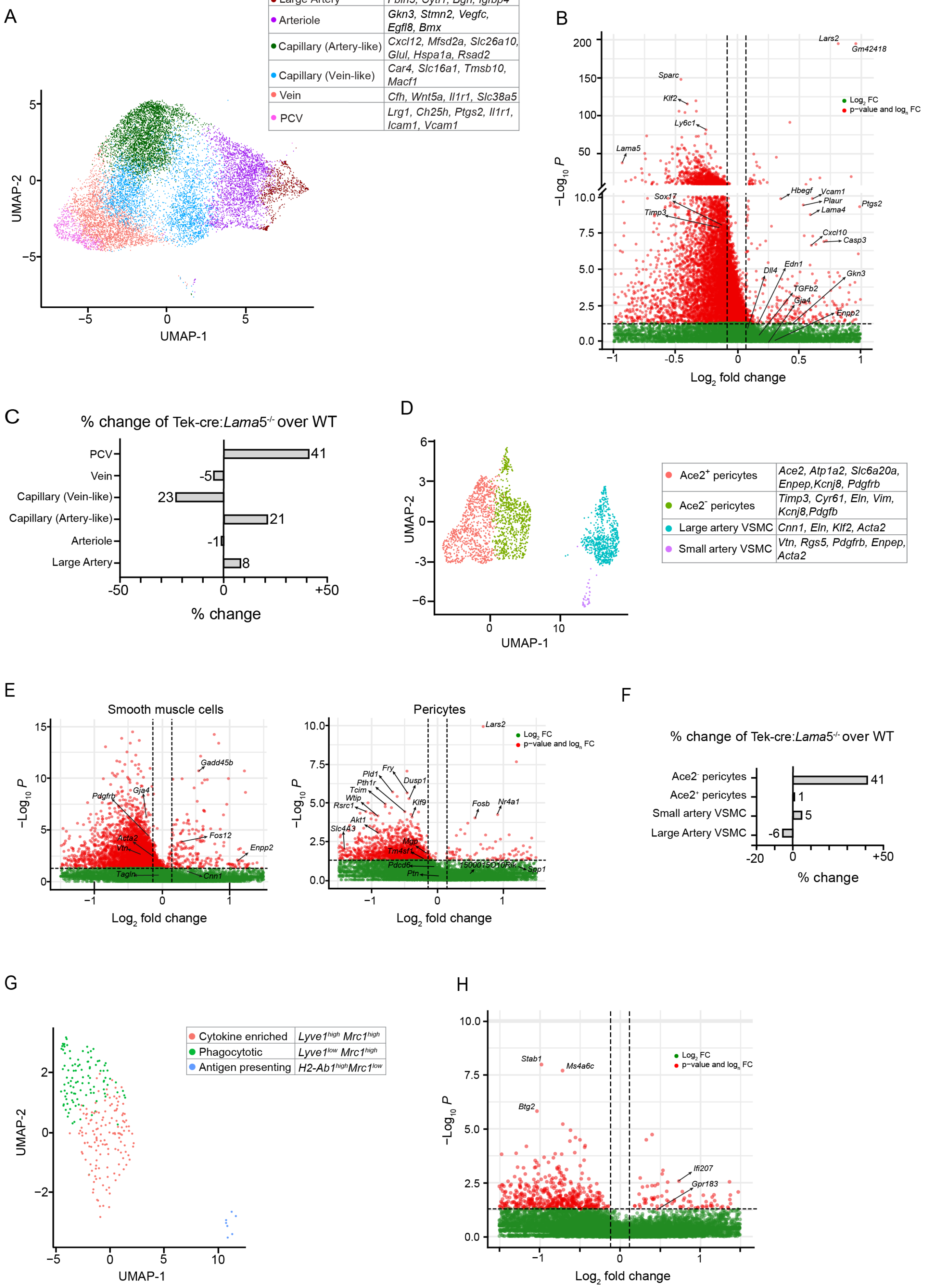


**sFigure 3 scRNAseq data for (A-C) endothelial cell populations, (D-F) smooth muscle cells and pericytes and (G, H) resident immune cells in enriched cerebral vessel samples from *Tek-Cre:Lama5*$^{-/-}$ and WT littermates.** UMAPs of **(A)** 6 endothelial **(D)** 2 smooth muscle and 2 pericyte, and **(G)** 3 myeloid cell populations identified. Different cells are colour coded based on their cluster affiliations shown in associated tables. Vulcano plots of genes upregulated and downregulation in *Tek-Cre:Lama5*$^{-/-}$ samples compared to WT littermates in **(B)** endothelial cells, **(E)** smooth muscle cells and pericytes and **(H)** myeloid cells; **(C,F)** corresponding graphs of percentage changes in *Tek-Cre:Lama5*$^{-/-}$ over WT littermates based on transcriptomic profiles.

Suppl. Figure 4

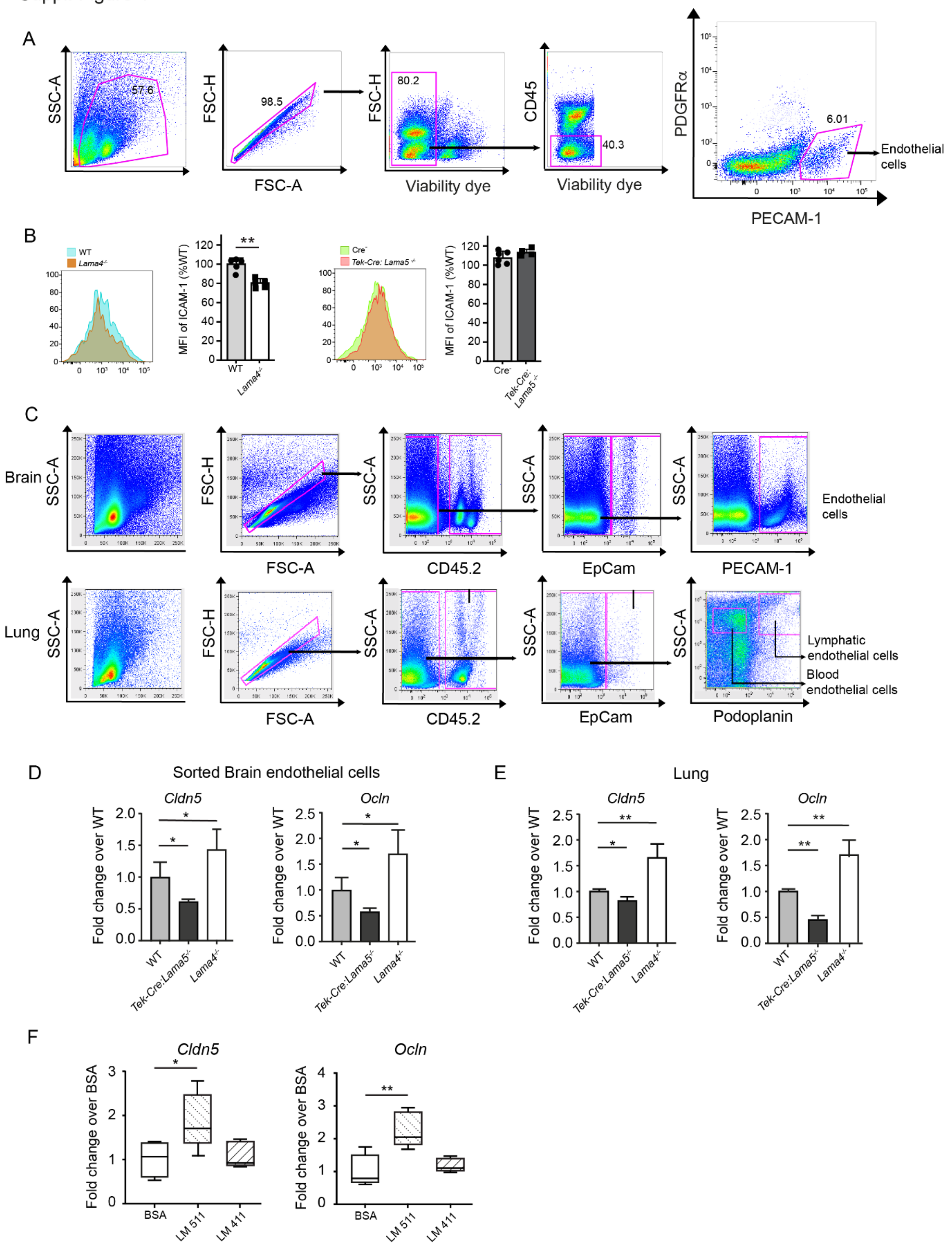


**sFigure 4 Analyes of tight junction molecules in brain and lung endothelium of naïve *Lama4*$^{-/-}$, *Tek-cre:Lama5*$^{-/-}$ and WT control mice.** (**A**) Flow cytometry strategy to identify PECAM$^{+}$ brain blood endothelial cells to determine ICAM-1 expression. **(B)** Representative FACS analyses of *Lama4*$^{-/-}$, *Tek-Cre:Lama5*$^{-/-}$ and WT control brain

samples for ICAM-1 and corresponding quantification (Unpaired t-Test, 5 mice per genotype in at least 2 separate experiments). (**C**) Flow cytometry strategy to distinguish brain and lung blood endothelial cells from podoplanin$^{+}$ lymphatic endothelial cells, EpCam$^{+}$ epithelial cells and CD45$^{+}$ leukocytes, which were employed for sorting of PECAM-1$^{+}$podoplanin$^{neg}$ blood endothelial cells. (**D,E**) qPCR for expression of occludin and claudin 5 in PECAM-1$^{+}$podoplanin$^{neg}$ blood endothelial cells from *Lama4*$^{-/-}$, *Tek-cre:Lama5*$^{-/-}$ and WT control (**D**) brains and (**E**) lungs; (ANOVA, 3 brains per genotype). (**F**) Quantification of *Cldn5* and *Ocln* expression in brain derived endothelioma cell line (bEND.5) plated on laminin 411, 511 or on BSA control, (ANOVA, triplicates per condition and two experiments). Data are means ± SD *$p$<0.05, **$p$<0.01, ***$p$<0.001.

Suppl. Figure 5

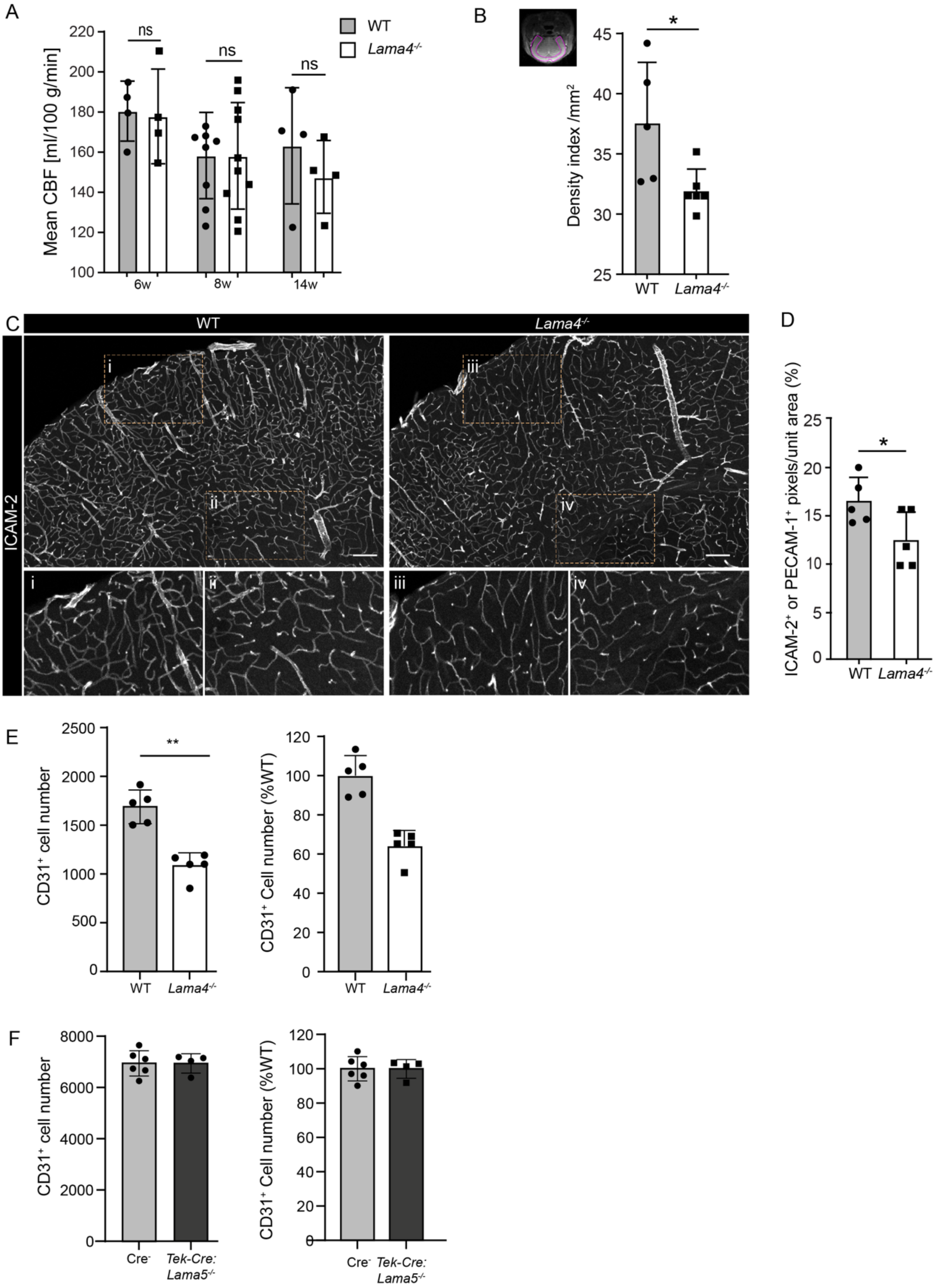


**sFigure 5 MRI and measures of vessel density.** MRI was performed to determine the (**A**) mean cerebral blood flow (CBF) and (**B**) microvessel density index (MDI) in the ROI (outlined in purple) of 14 week-old naïve *Lama4*$^{-/-}$ and WT control mice; (unpaired t-Test; at least 4 mice per genotype). (**C**) Representative

immunofluorescence staining for ICAM-2 to mark all blood vessels in *Lama4*$^{-/-}$ and WT control mice and (**D**) corresponding quantification of pixels/unit area using ICAM-2 (2 mice) or CD31 (3 mice) immunofluorescent staining (unpaired t-Test; at least 5 mice per genotype in 2 experiments). (**E** and **F**) Flow cytometry quantification of endothelial cells in *Lama4*$^{-/-}$, *Tek-Cre:Lama5*$^{-/-}$ and WT control brains (unpaired t-Test, 4-6 brains per genotype were analysed in at least 2 experiments). Data are means ± SD *$p$<0.05, **$p$<0.01, ***$p$<0.001. Scale bar in C = 100 µm

Suppl. Figure 6

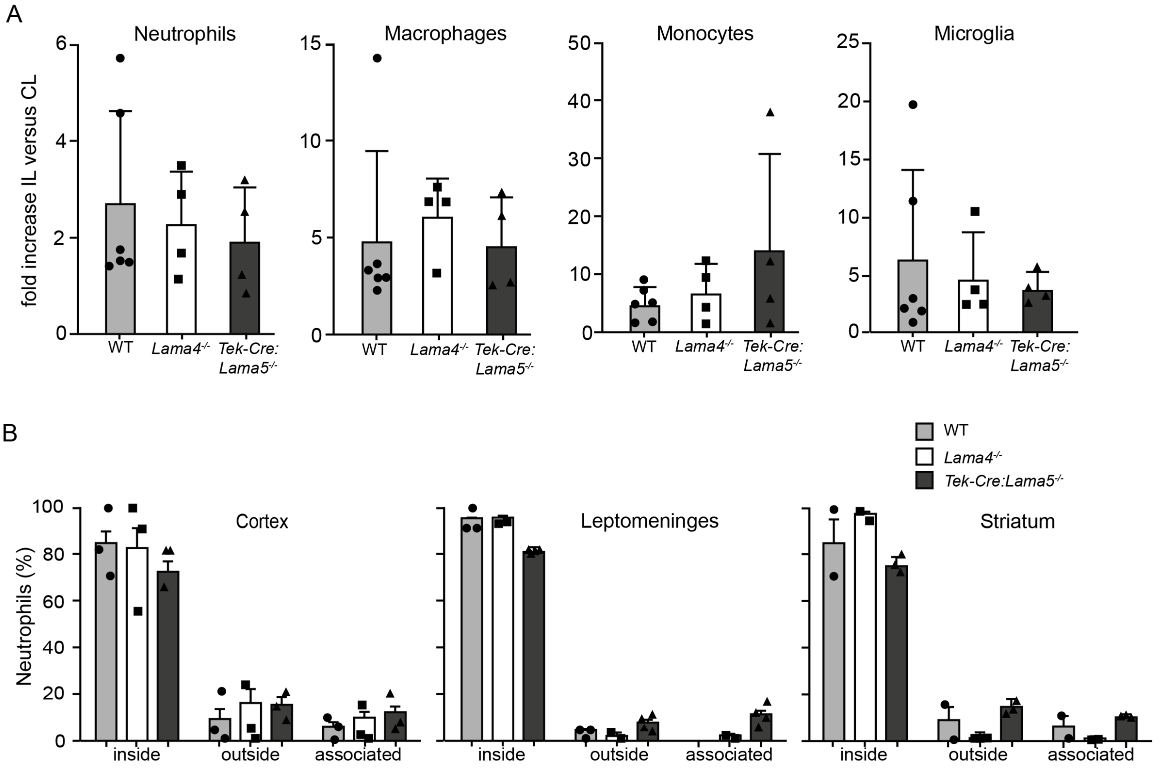


**sFigure 6 Immune cell infiltration after stroke (60min tMCAO/24h reperfusion) in *Lama4*$^{-/-}$, Tek-cre:*Lama5*$^{-/-}$ and WT control mice.** (**A**) Flow cytometry quantification of Ly6G$^{+}$ neutrophils, F4/80$^{+}$ macrophages, Ly6C$^{+}$ monocytes and Iba$^{+}$ microglia in ipsilateral (IL) and contralateral (CL) hemispheres of *Lama4*$^{-/-}$, Tek-cre:*Lama5*$^{-/-}$ and WT control mice; data are expressed as fold increase in cell numbers in IL/CL hemispheres (ANOVA followed by post hoc Tukey, 4-6 brains per genotype from at least 3 experiments). (**B**) Quantification of Ly6G$^{+}$ neutrophils within vessel borders, determined by pan-laminin immunofluorescence staining, outside of vessels or associated with vessel walls [4] (ANOVA followed by post hoc Tukey, 4 brains per genotype using 3 sections (100 µm thick) for each brain and 5 images for each area (leptomeninges, cortex and striatum)). Data are means ± SD.

Suppl. Figure 7

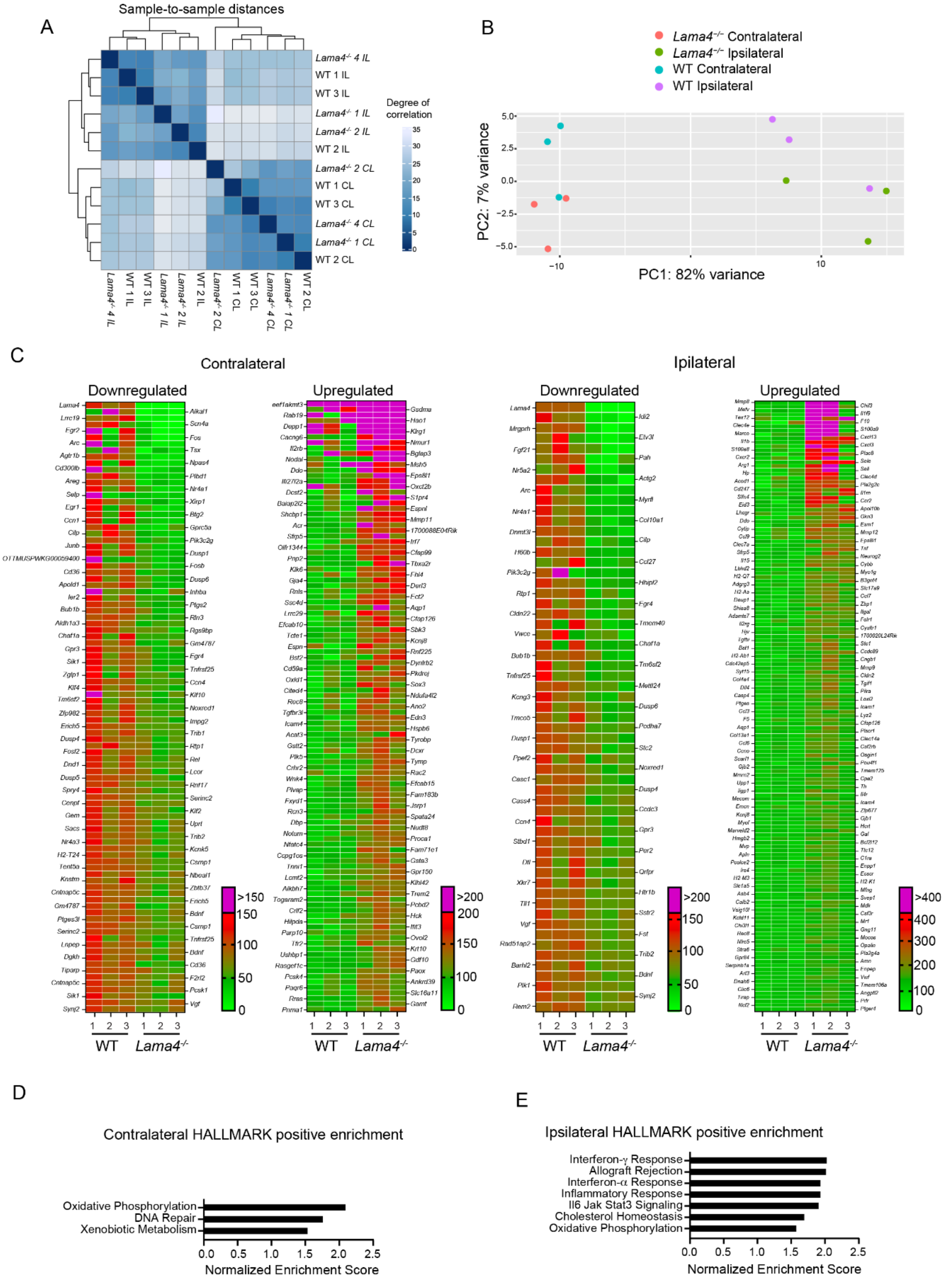


**sFigure 7 Bulk RNA sequencing data from IL and CL brain hemispheres of *Lama4*$^{-/-}$ and WT control mice after stroke (60min tMCAO/24h reperfusion).**

(**A**) Heatmap of the sample-to-sample distances based on the count data for gene expression. The hierarchical clustering shows the closely related samples being grouped together. The colour intensity indicates strength of sample-to-sample association. (**B**) Principal component analysis showing differences between samples. (**C**) Heatmap of genes with $p \leq 0.01$ for IL and CL hemispheres of WT and *Lama4*$^{-/-}$ mice; rows represent genes and each column represents one mouse; colour coding represents the mean expression (transcripts per million). Gene set enrichment analysis (GSEA) of contralateral (**D**) and ipsilateral (**E**) brain hemispheres shows the enriched gene sets/pathways from HALLMARK data bases.

Suppl. Figure 8

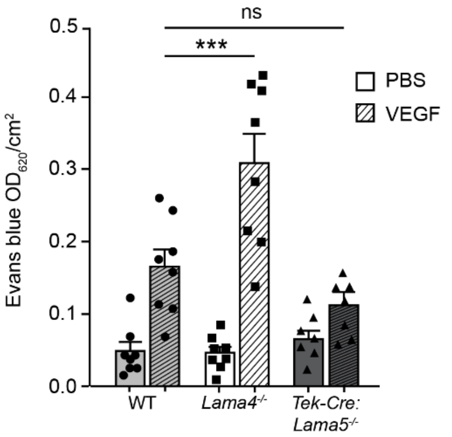


**sFigure 8 Modified Miles permeability assay.**

Extravasated Evans Blue in the dermis of *Lama4*$^{-/-}$, *Tek-Cre:Lama5*$^{-/-}$ and WT naïve mice was measured at $OD_{620}$ in response to VEGF or PBS injection (2way-ANOVA, 7-8 mice per genotype in 3 experiments). Data are means ± SD *$p<0.05$, **$p<0.01$, ***$p<0.001$.

**Supplementary Table 1. Primary Antibodies Employed in Immunofluorescence Analyses (IF), Western Blots (WB) and Flow Cytometry (FC).**

| Target Molecule | Antibody Designation/Clone | Application | Reference/Source |
|---|---|---|---|
| ACE-2 | goat anti-mouse | IF | R&D Systems/AF3437 |
| CD11b APC | rat anti-mouse/M1/70 | FC | eBioscience 17-0112-83 |
| CD102 (ICAM-2) | rat anti-mouse/UZ10* | IF | RH. Personal communication |
| CD106 (VCAM-1) | rat anti-mouse/M/K-2 | IF | SouthernBiotech 1510-01 |
| CD140a (PDGFRα) PE | rat anti-mouse/APA5 | FC | Thermo Fisher 12-1401-81 |
| CD140b (PDGFRβ) | Rat anti-mouse/APB5 | IF | Thermo Fisher 16-1402-82 |
| CD206 | Goat anti-mouse | IF | R&D Systems/AF2535 |
| | Goat anti-mouse | WB | R&D Systems/AF643 |
| CD31 (PECAM-1) | rat anti-mouse/MEC 13.3 | IF, WB | BD Pharmingen/553370 |
| | Hamster anti-mouse/2H8 | IF, WB | Abcam/ab119341 or Thermo Fisher MA3105 |
| CD31 (PECAM-1) FITC | rat anti-mouse/MEC13.3 | FC | BD bioscience 551262 |
| | rat anti- mouse/390 | FC | Biolegend 102406 |
| CD326 (EpCam) | rat anti-mouse/G8.8 | FC | BD bioscience 565425 |
| CD45 | rat anti-mouse/30G12 | IF | [5] |
| CD45.2 PE | mouse anti-mouse/104 | FC | eBioscience 12-0454-83 |
| CD45.2 PerCP/cy5.5 | mouse anti-mouse/104 | FC | Biolegend 109828 |
| CD54 (ICAM-1) | goat anti-mouse | WB | R&D Systems/AF796-SP |
| CD54 (ICAM-1) BV421 | hamster anti-mouse/3E2 | FC | BD Biosciences 752967 |
| Claudin 5 | rabbit anti-mouse | IF, WB | Invitrogen/34-1600 |
| Connexin 37 | rabbit anti-mouse | IF | Thermo Fisher/40-4300 |
| | rabbit anti-mouse | IF | Thermo Fisher/42-4400 |
| Dll4 | goat anti-mouse | IF | R&D Systems/AF1389 |
| ENPP2/ATX | mouse anti-human | IF | Abcam/ab77104 |
| | rabbit ant-human | IF | Abcam/ab137590 |
| | rabbit anti-human | WB | Proteintech/14243-1-AP |
| GAPDH | rabbit anti-human/14C10 | WB | Cell Signalling/2118 |
| IgG | Cy3 goat anti-mouse | IF | Dianova/115-165-062 |
| Laminin α4 | Rabbit anti-mouse/377 | IF | [6] |
| Laminin α5 | Rat anti-mouse/4G6 | IF | [7] |
| Ly6C PerCP/Cy5.5 | rat anti-mouse/HK1.4 | FC | eBioscience 45-5932-82 |

| | | | |
|---|---|---|---|
| Ly6G FITC | rat anti-mouse/1A8 | FC | Bio-Legend 127606 |
| MHCII | Rat anti-mouse | IF | Serotec/MCA949F |
| Occludin | Rabbit anti human | IF | Abcam 31721 |
| | Rabbit anti human | IF | Thermo Scientific 71-1500 |
| | Rabbit anti human | WB | Thermo Fisher/40-4700 |
| Pan Laminin | rabbit anti-mouse (ab455) | IF | [8] |
| Podoplanin | hamster anti-mouse/8,1 | FC | eBioscience 25-5381-803 |
| PTGIS | rabbit anti-human | WB | Proteintech/27061-1-AP |
| SMA | Mouse anti-mouse/1A4 | IF | Sigma/C6198 |
| | Rabbit anti-human | IF, WB | Merck/Millipore/PA5-16697 |

#### Supplementary Table 2. Secondary Antibodies Employed

| **Antigen** | **Fluorophor** | **Application** | **Dilution/ concentration** | **Company/ Catalogue No.** |
|---|---|---|---|---|
| donkey anti-goat | Alexa Fluor 488 | IF | 1:400/5 µg\|ml | Molecular Probes/A11055 |
| donkey anti-goat | Alexa Fluor 488 | IF | 1:1000/2 µg/ml | Abcam/ab150133 |
| donkey anti-rabbit | Alexa Fluor 568 | IF | 1:400/5 µg\|ml | Abcam/175692 |
| donkey anti-rabbit | Alexa Fluor 647 | IF | 1:1000/2 µg/ml | Abcam/150067 |
| donkey anti-rabbit | Alexa Fluor 488 | IF | 1:1000/2 µg/ml | Abcam/ab 150065 |
| donkey anti-rabbit | Alexa Fluor 594 | IF | 1:2000/0.25 µg/ml | Invitrogen/A21207 |
| donkey anti-rat | Alexa Fluor 647 | IF | 1:1000/2 µg/ml | Abcam/ab 150155 |
| goat anti-mouse | Cy3 | IF | 1:800/1.8 µg/ml | Dianova/115-165-062 |
| goat anti-rat | Alexa Fluor 488 | IF | 1:2000/1 µg/ml | Molecular Probes/A11006 |
| donkey anti-sheep/goat | HRP | WB | 1:5000 | Millipore/AB324P |
| goat anti-rabbit | HRP | WB | 1:5000 | Bio-Rad/ 172-1019 |
| viability dye | eFluor 780 | FC | 1:1000 | Biosciences 65-0865-14 |

#### Supplementary Table 3. Full names of the gene symbols mentioned in the text

| Gene Symbol | Full Name |
|---|---|
| *Acta2* | Actin, alpha 2, smooth muscle, aorta |
| *Atf3* | Activating transcription factor 3 |
| *Ccl3* | Chemokine (C-C motif) ligand 3 |
| *Ccl7* | Chemokine (C-C motif) ligand 7 |
| *Ccr2* | C-C chemokine receptor type 2 |
| *Cd68* | CD68 molecule (macrosialin) |
| *Chil3* | Chitinase-like 3 |
| *Cldn5* | Claudin 5 |
| *Cnn1* | Calponin 1 |
| *Ctsd* | Cathepsin D |
| *Cxcl13* | Chemokine (C-X-C motif) ligand 13 |
| *Dll4* | Delta like canonical Notch ligand 4 |
| *Edn1* | Endothelin 1 |
| *Ednrb* | Endothelin receptor type B |
| *Enpp2* | Ectonucleotide pyrophosphatase/phosphodiesterase 2 |
| *Fbln5* | Fibulin 5 |
| *Gja1* | Gap junction protein, alpha 1 |
| *Gskn3* | Unconfirmed symbol (possible typo, maybe Gsk3b?) |
| *H2-ab1* | Histocompatibility 2, class II antigen A, beta 1 |
| *Icam1* | Intercellular adhesion molecule 1 |
| *Klf2* | Kruppel-like factor 2 |
| *Lama4* | Laminin subunit alpha 4 |
| *Lama5* | Laminin subunit alpha 5 |
| *Lyve1* | Lymphatic vessel endothelial hyaluronan receptor 1 |
| *Marco* | Macrophage receptor with collagenous structure |
| *Mmp8* | Matrix metallopeptidase 8 |
| *Mrc1* | Mannose receptor, C-type 1 |
| *Ms4a7* | Membrane spanning 4-domains A7 |
| *Ocln* | Occludin |
| *Pf4* | Platelet factor 4 |
| *Ptgis* | Prostaglandin I2 (prostacyclin) synthase |
| *Tagln* | Transgelin |
| *Tek* | TEK receptor tyrosine kinase (Tie2) |
| *Tgfb2* | Transforming growth factor, beta 2 |
| *Ttr* | Transthyretin |
| *Vcam1* | Vascular cell adhesion molecule 1 |
| *Vtn* | Vitronectin |
| *Wfdc12* | WAP four-disulfide core domain 12 |
| *Wfdc17* | WAP four-disulfide core domain 17 (not fully confirmed) |

#### Supplementary Table 4. Primer sets used in qPCR

| Gene/protein | Forward | Reverse |
|---|---|---|
| *Cldn5*/ claudin-5 | 5’ GCAAGGTGTATGAATCTGTGCT-3’ | 5’-GTCAAGGTAACAAAGAGTGCCA-3’ |
| *Gapdh*/GAPDH | 5’-TGGCCTTCCGTGTTCCTAC-3’ | 5’-GAGTTGCTGTTGAAGTCGCA-3’ |
| *Ocln*/occludin | 5’-TTGAAAGTCCACCTCCTTACAGA-3’ | 5’-CCGGATAAAAAGAGTACGCTGG-3’ |
